# Iro-C/IRX creates anti-cancerized epithelial field against IL-6-dependent malignant tumorigenesis

**DOI:** 10.64898/2026.01.21.700777

**Authors:** Tomonori Nakanishi, Masato Enomoto, Tatsushi Igaki

**Affiliations:** Laboratory of Genetics, Graduate School of Pharmaceutical Sciences, Kyoto University, Yoshida-Shimoadachi-cho, Sakyo-ku, Kyoto 606-8501, Japan; Laboratory of Genetics, Graduate School of Biostudies, Kyoto University, Yoshida-Shimoadachi-cho, Sakyo-ku, Kyoto 606-8501, Japan; Laboratory of Dynamic Biology, Department of Integrative Vascular Biology, Faculty of Medical Sciences, University of Fukui, 23-3, Matsuoka-shimoaizuki, Eiheiji-cho, Yoshida-gun, Fukui, 910-1193, Japan

## Abstract

Some epithelial regions appear intrinsically resistant to malignancy, yet how such anti-cancerized fields are established remains elusive. Here, we show in *Drosophila* that Iroquois Complex (Iro-C), a group of transcriptional repressors specifically expressed in the notum region of the wing imaginal epithelium, creates anti-cancerized field against malignant tumorigenesis. Clones of cells with Ras activation and cell polarity defect (Ras^V12^*/scrib^-/-^*) develop into malignant tumors in the pouch and hinge regions of the wing disc, yet they failed to overgrow in the notum. Mechanistically, intrinsic Iro-C expression in the notum represses *upd/IL-6* transcription, thereby preventing JAK-STAT activation essential for driving malignant growth. Forced expression of Upd/IL-6 converted the notum into tumor-prone field, whereas forced expression of Iro-C in the pouch and hinge abolished their Ras^V12^*/scrib^-/-^* tumorigenesis. Our findings provide a mechanistic explanation for how some epithelial regions, such as distal segments of the kidney, where Iro-C/IRX1 is specifically expressed, exhibit resistance to cancer development, and offer a novel therapeutic strategy against IL-6-dependent cancers.

## Introduction

Cancer development within a given organ is not uniform; rather, it exhibits striking regional variation shaped by both intrinsic cellular properties and the surrounding tissue microenvironment. For instance, breast cancers predominantly arise in the upper-outer quadrant of the mammary gland, while other regions exhibit lower tumor incidence (Lee, 2005). In the kidney, renal cancers commonly originate from proximal tubule cells and intercalated cells of the collecting duct, whereas cells in distal segment are rarely affected (Lindgren *et al*, 2017). Colorectal cancers more frequently occur in the left-sided colon (descending colon, sigmoid colon, and rectum) than in the right-sided colon (cecum and ascending colon) (Ikuta *et al*, 2024; Topdagi & Timuroglu, 2018). Furthermore, pancreatic cancer frequently arises in the head of the pancreas, whereas body/tail tumors are less common (Luo *et al*, 2020). While most studies have focused on elucidating why tumors preferentially develop in certain cancer-prone regions or cell populations, much less is known about how certain regions within a tissue actively resist oncogenic transformation.

Studies in *Drosophila* have provided important insights into how local tissue context influences tumor behavior. In *Drosophila* wing imaginal epithelium, cells deficient for neoplastic tumor suppressor genes (*nTSG*) develop into tumors in the hinge region, where the endogenous JAK-STAT activity is high and the cytoskeletal organization is distinct from other regions with basally enriched microtubules (Tamori *et al*, 2016). Intriguingly, however, *nTSG*-deficient cells do not overgrow but are eliminated in the pouch region of the wing imaginal epithelium through cell competition (Agrawal *et al*, 1995; Brumby & Richardson, 2003; Igaki *et al*, 2009; Woods & Bryant, 1991). In addition, cells deficient for *polyhomeotic* (*ph*), a component of the Polycomb repressive complex (PRC1), overgrow in the pouch and hinge regions but not in the notum region of the wing disc. This growth restriction was attributed to the lower responsiveness of notum cells to JNK and JAK-STAT signaling (Medina *et al*, 2021). However, the molecular basis that actively confers the tumor resistance on notum cells remains unexplored.

Here, using the flippase (FLP)-flippase recombination target (FRT)-mediated genetic mosaic technique (Lee & Luo, 1999; Xu & Rubin, 1993), we show in *Drosophila* wing imaginal epithelia that Ras-activated malignant tumors, but not hyperplastic benign tumors, exhibit strong regional preference to overgrow and uncover the underlying molecular underpinnings of how such anti-cancerized field is created.

## Results

### The notum region behaves as anti-cancerized field against malignant tumorigenesis

In *Drosophila* wing imaginal epithelium (Figure 1A), cell clones expressing oncogenic Ras (Ras^V12^) exhibit hyperproliferation compared to wild-type clones, as previously reported (Brumby & Richardson, 2003; Igaki *et al*, 2006; Pagliarini & Xu, 2003) (Figure 1D, compare to 1B, quantified in 1G and 1I, respectively). In contrast, cell clones lacking apico-basal polarity gene, such as *scribble* (*scrib*) gene, are eliminated from the tissue by cell competition (Brumby & Richardson, 2003; Igaki *et al*., 2009) (Figure 1C, quantified in 1H). However, when *scrib* loss is combined with Ras activation (Ras^V12^*/scrib^-/-^*), the resulting cell clones exhibit malignant tumorigenesis characterized by invasive overgrowth (Brumby & Richardson, 2003; Pagliarini & Xu, 2003). Intriguingly, while Ras^V12^*/scrib^-/-^* clones displayed extensive overgrowth in the pouch and hinge (P+H) regions of the wing disc (Figure 1A and 1E), growth of these clones was significantly suppressed in the notum (N) region (Figure 1E, quantified in 1J, P+H/N boundary shown by an yellow dotted line in Figure 1F). A similar regional tumor suppression was observed when clones of cells with Ras activation and *scrib*-knockdown (Ras^V12^*/scrib*.IR) were induced in the wing disc (Figure S1A, quantified in S1B). Moreover, induction of Ras^V12^*/scrib*.IR cells in the pouch region by *nub*-Gal4 resulted in aggressive overgrowth (Figure S1C and S1D, quantified in S1G), whereas Ras^V12^*/scrib*.IR cells induced in the notum region by *76A01*-Gal4 did not show growth advantage compared to the control (Figure S1E and S1F, quantified in S1G). To determine the developmental stage at which clones begin to exhibit differential growth between the notum and P+H regions, we performed a time-course analysis using a heat-shock-induced MARCM system to induce Ras^V12^/*lgl^-/-^* clones. At 89-97 h AEL, the relative size of Ras^V12^/*lgl^-/^* clones was nearly identical between the notum and P+H regions (Figure S1H, quantified in S1L). However, at later stages (113-121 h AEL and 137-145 h AEL), clones in the P+H region showed robust overgrowth, whereas those in the notum region do not increase further in size (Figure S1I and S1J, quantified in S1K and S1L). In contrast, wild-type clones showed no difference in size between the notum and P+H regions at 89-97 h AEL and 113-121 h AEL (Figure S1M and S1N, quantified in S1O and S1P); wild-type larvae had already pupariated by 137-145 h AEL. These data indicate that the differential growth phenotype between the two regions emerges between 97 and 113 h AEL.

**Figure 1.**
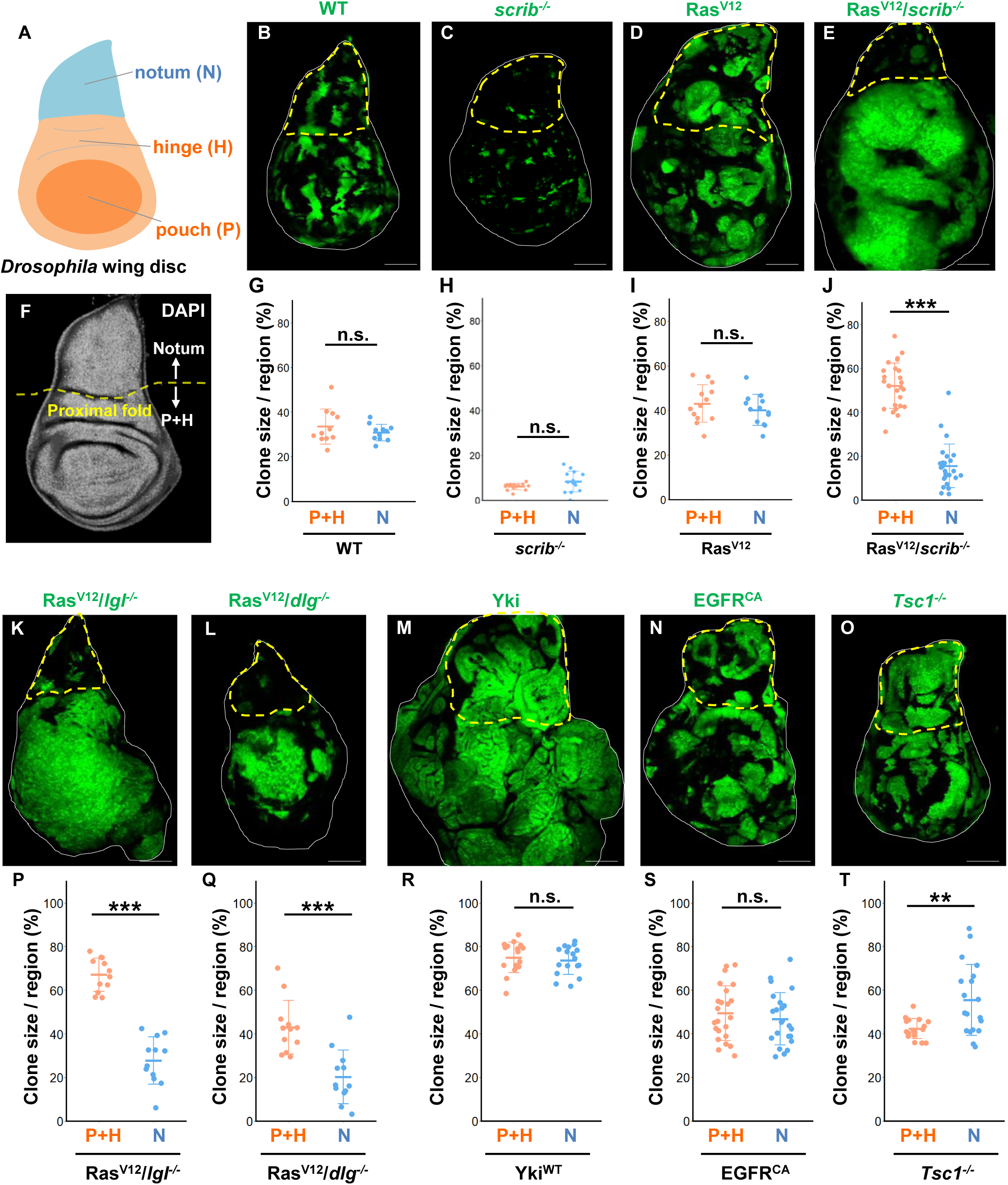
The notum region behaves as anti-cancerized field against malignant tumorigenesis. (A) A schematic Figure showing 3 subdomains of a third instar wing imaginal disc; pouch, hinge, and notum. (B-E) Wing discs bearing GFP-labeled wild-type (B), *scrib^-/-^* (C), Ras^V12^ (D), and Ras^V12^/*scrib^-/-^* (E) clones, dissected at 96-120 hour after egg lying (96-120 h AEL) (B) and (C), 120-144 h AEL (D) or 144-168 h AEL (E). (F) Wing disc bearing GFP-labeled wild-type clones, stained with DAPI. The proximal fold structure between the notum and hinge regions is indicated by yellow dotted lines. (G-J) Quantification of the clone size in the Pouch+Hinge (P+H) or Notum (N) (% of the clone area per region area) for wild-type (n=11, p=0.5545), *scrib^-/-^* (n=14, p=0.1029), Ras^V12^ (n=13, p=0.3358), and Ras^V12^/*scrib^-/-^* (n=25, p<0.0001) clones. ***p < 0.001; n.s. (not significant); Wilcoxon rank sum test. (K-O) Wing discs bearing GFP-labeled Ras^V12^/*lgl^-/-^* (K) and Ras^V12^/ *dlg^-/-^* (L), Yki (M), EGFR^CA^ (N), and *Tsc1^-/-^* (O) clones, dissected at 120-144 AEL (M), (N), and (O), 144-168 h AEL (K) and (L). (P-T) Quantification of the clone size in the Pouch+Hinge (P+H) or Notum (N) (% of the clone area per region area) for Ras^V12^/*lgl^-/-^* (n=12, p<0.0001), Ras^V12^/*dlg^-/-^*(n=12, p=0.0003), Yki (n=17, p=0.5177), EGFR^CA^ (n=23, p=0.4551), and *Tsc1^-/-^* (n=19, P=0.0051) clones. ***p < 0.001; **p < 0.01; n.s. (not significant); Wilcoxon rank sum test. Scale bar, 100 µm.

Furthermore, this regional tumor suppression was also observed in other Ras-activated neoplastic tumor clones, such as Ras^V12^*/lgl^-/-^*and Ras^V12^*/dlg^-/-^* tumors (Igaki *et al*., 2006) (Figure 1K and 1L, quantified in 1P and 1Q). In contrast, hyperplastic tumor models, including Ras^V12^ clones, Yorkie (Yki; a YAP homolog)-overexpressing clones, EGFR-activated (EGFR^CA^) clones, and *Tsc1* mutant (*Tsc1^-/-^*) clones, uniformly overgrew throughout the wing disc, including the pouch, hinge, and notum regions (Figure 1D and 1M-1O, quantified in 1H and 1R-1T). These observations suggest that the notum region of the wing imaginal epithelium behaves as anti-cancerized field that specifically prevents Ras-activated malignant tumorigenesis.

### The notum region suppresses proliferative activity of Ras^V12^/*scrib*^-/-^ tumors

To dissect the mechanism underlying notum-specific tumor suppression, we first assessed the role of cell death, a known mediator of intrinsic tumor suppression (Kanda, 2020; Lowe *et al*, 2004). Detection of dying cells by the caspase activity probe CD8-PARP-Venus combined with anti-cleaved-PARP staining revealed no significant difference in the number of cell death in Ras^V12^*/scrib*.IR cells between the notum and P+H regions (Figure 2A and 2B, quantified in 2C). In addition, genetic inhibition of apoptosis by *miRHG* (miRNA targeting three pro-apoptotic genes *reaper* (*rpr*), *hid*, and *grim*) in Ras^V12^*/scrib^-/-^* clones did not rescue their proliferation in the notum (Figure 2D and 2E, quantified in 2F), indicating that increased cell death is not the mechanism for tumor suppression in this region. We next examined cell proliferation using phospho-histone H3 (PH3) as a mitosis marker and EdU incorporation for DNA synthesis. Notably, the number of PH3- and EdU-positive cells was significantly lower in Ras^V12^*/scrib^-/-^* clones located in the notum compared to the P+H regions (Figure 2G-2J, quantified in 2O and 2P). Importantly, in wild-type tissue, proliferative activity in the notum was comparable to that in the P+H regions (Figure 2K-2N, quantified in 2O and 2P). Together, these results suggest that the notum region specifically suppresses proliferative activity of Ras^V12^*/scrib^-/-^* tumors.

**Figure 2.**
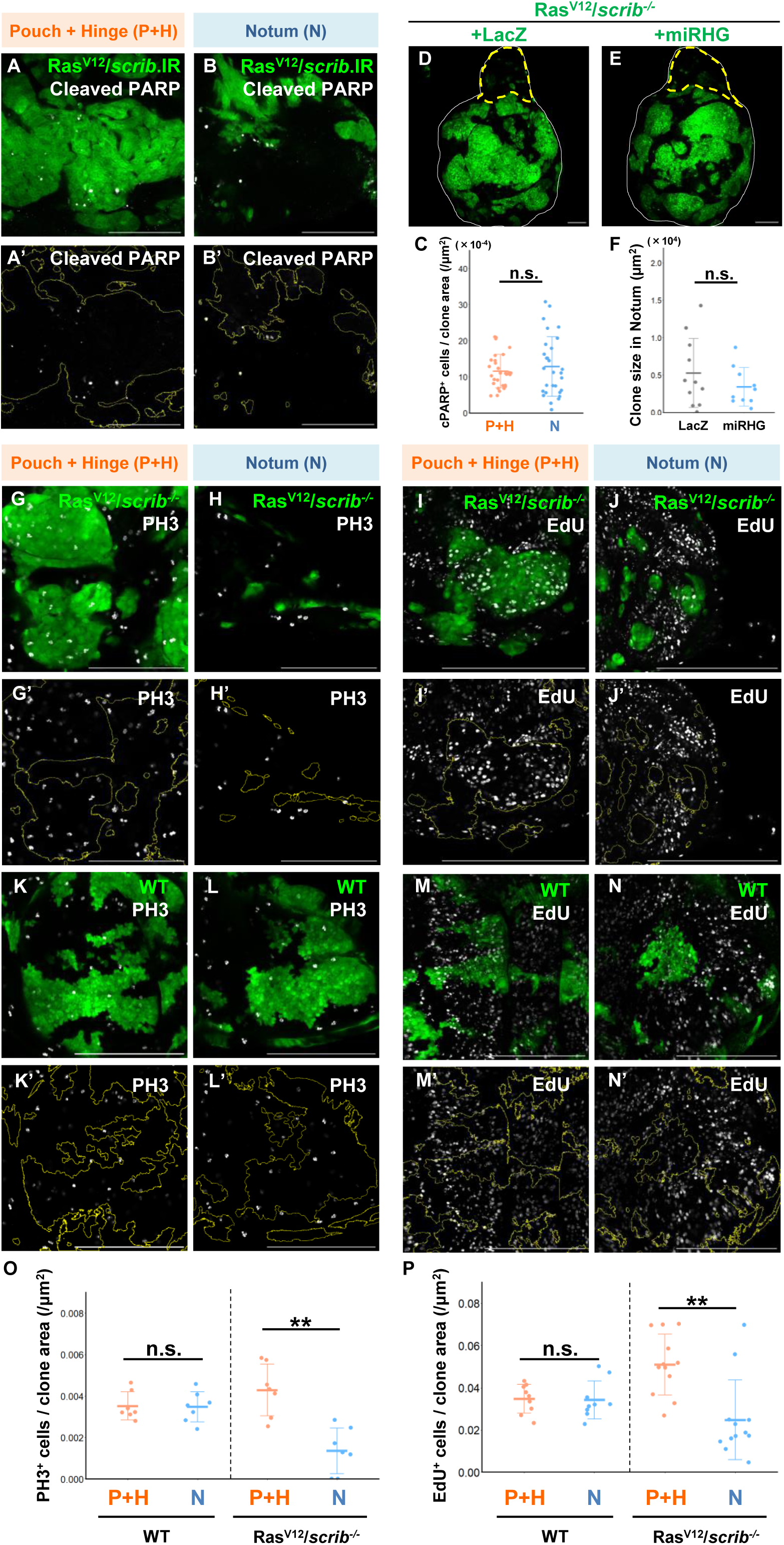
The notum region suppresses proliferative activity of Ras^V12^/*scrib*^-/-^tumors. (A and B) Pouch+Hinge (P+H) region (A) or Notum (N) region (B) of the wing disc bearing GFP-labeled Ras^V12^+*scrib.IR* clones stained with anti-cleaved PARP antibody, dissected at 144-168 h AEL. (C) Quantification of the number of dying cells (cPARP-positive cells) in the Ras^V12^+*scrib.IR* clones in the P+H or N region (n=29, p=0.9133). n.s. (not significant); Wilcoxon rank sum test. (D and E) Wing discs bearing GFP-labeled Ras^V12^/*scrib^-/-^*+LacZ (D) and Ras^V12^/*scrib^-/-^*+miRHG (E) expressing clones, dissected at 144-168 h AEL. (F) Quantification of the clone size in the notum region for Ras^V12^/*scrib^-/-^*+LacZ (n=11) and Ras^V12^/*scrib^-/-^*+miRHG (n=10, p=0.7564) expressing clones. n.s. (not significant); Wilcoxon rank sum test. (G, H, K, and L) Pouch+Hinge (P+H) region (G and K) or Notum (N) region (H and L) of the wing disc bearing GFP-labeled wild-type or Ras^V12^/*scrib^-/-^* clones stained with anti-PH3 antibody, dissected at 120-144 h AEL (wild-type) or 144-168 h AEL (Ras^V12^/*scrib^-/-^*). (I, J, M, and N) Pouch+Hinge (P+H) region (I and M) or Notum (N) region (J and N) of the wing disc bearing GFP-labeled wild-type or Ras^V12^/*scrib^-/-^* clones stained with EdU, dissected at 120-144 h AEL. (O) Quantification of the PH3-positive cells in wild-type (n=7, p=0.9015) or Ras^V12^/*scrib^-/-^* (n=7, p=0.0033) clones in the P+H or N region. **p < 0.01; n.s. (not significant); Wilcoxon rank sum test. (P) Quantification of the EdU-positive cells in wild-type (n=9, p=0.6588) or Ras^V12^/*scrib^-/-^* (n=12, p=0.0020) clones in the P+H or N region. **p < 0.01; n.s. (not significant); Wilcoxon rank sum test. Scale bar, 100 µm.

### Notum-specific expression of Iro-C represses *upd/IL-6* transcription

To uncover the factors responsible for creating anti-cancerized field in the notum, we screened a series of genes specifically expressed in the notum (Figure S2A). As a result, we found Iroquois Complex (Iro-C) genes—*araucan (ara)*, *mirror (mirr)*, and *caupolican (caup)*—as factors that confer tumor resistance on notum cells. Knockdown of Iro-C resulted in massive overgrowth of Ras^V12^*/scrib^-/-^*clones specifically in the notum (Figure 3A-3C, quantified in 3D), but not in the pouch and hinge regions (quantified in 3E), while Iro-C knockdown alone only mildly affected the clone size in the notum (Figure S2B-S2D, quantified in S2E). Iro-C genes encode evolutionary conserved homeodomain transcriptional repressors that are mainly expressed in the notum region of the wing disc (Cavodeassi *et al*, 2001; Villa-Cuesta & Modolell, 2005) (Figure S2F-S2H). The vertebrate homolog of Iro-C is IRX family proteins (IRX1∼IRX6), which also play key roles in cell specification in various tissues of *Xenopus*, zebrafish, mice, and humans (Alarcón *et al*, 2008; Bellefroid *et al*, 1998; Cavodeassi *et al*., 2001; Gómez-Skarmeta *et al*, 2001; Gómez-Skarmeta *et al*, 1998; Houweling *et al*, 2001; Kim *et al*, 2012; Marra & Wingert, 2014; McDonald *et al*, 2010).

**Figure 3.**
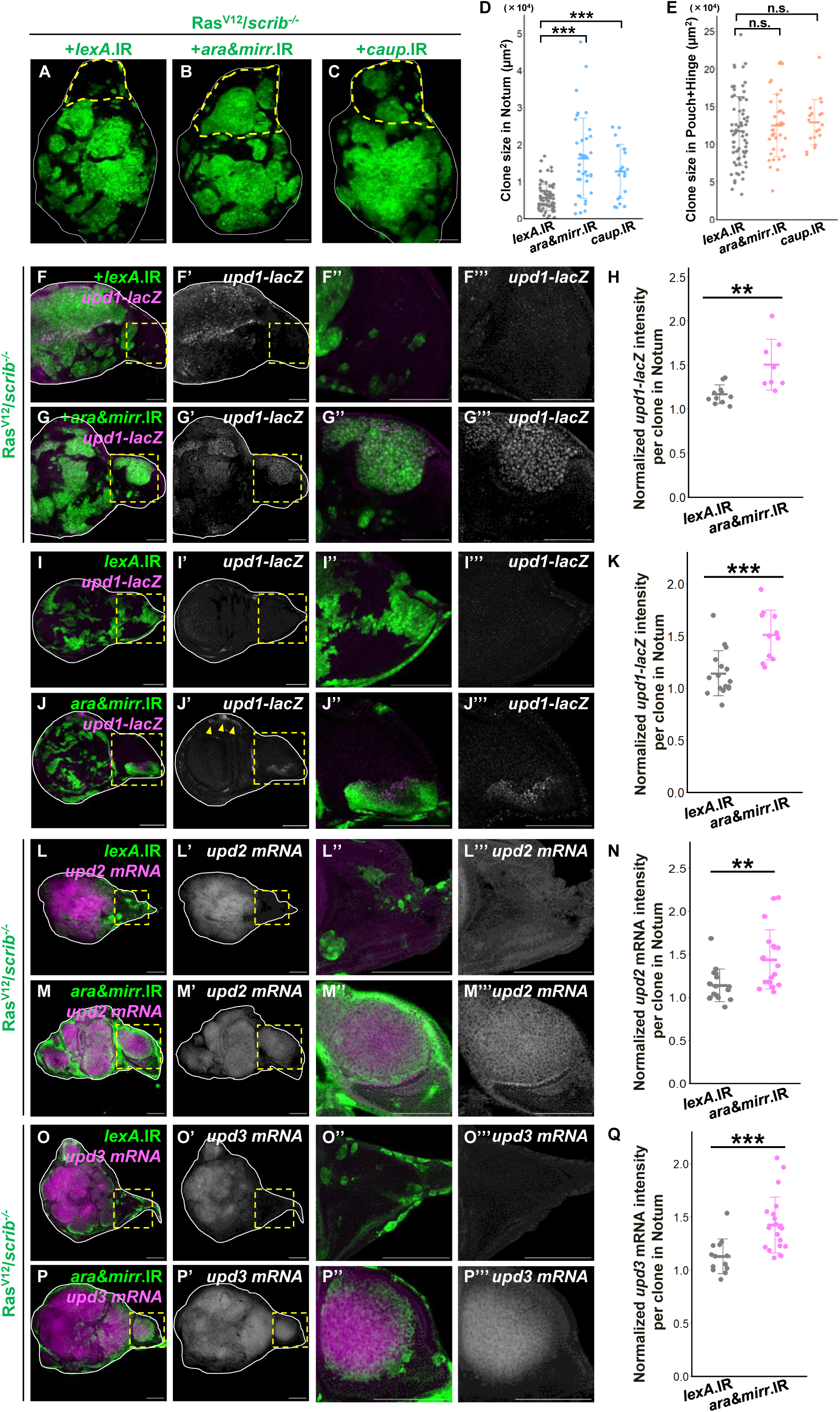
Notum-specific expression of Iro-C represses *upd/IL-6* transcription. (A-C) Wing discs bearing GFP-labeled Ras^V12^/*scrib^-/-^*+*lexA*.IR (A), Ras^V12^/*scrib^-/-^*+*ara&mirr.*IR (B), and Ras^V12^/*scrib^-/-^*+*caup.*IR (C) clones, dissected at 144-168 h AEL. (D) Quantifications of the clone size in the Notum region for Ras^V12^/*scrib^-/-^*+*lexA*.IR (n=71), Ras^V12^/*scrib^-/-^*+*ara&mirr.*IR (n=37, p<0.0001), and Ras^V12^/*scrib^-/-^*+*caup.*IR (n=20, p=0.0002) clones. ***p < 0.001; Kruskal-Wallis test followed by Dunn’s many-to-one comparison test with Holm correction. followed by Dunn’s many-to-one comparison test with Holm correction. (E) Quantifications of the clone size in the Pouch+Hinge (P+H) regions for Ras^V12^/*scrib^-/-^*+*lexA*.IR (n=71), Ras^V12^/*scrib^-/-^*+*ara&mirr.*IR (n=37, n=0.3487), and Ras^V12^/*scrib^-/-^*+*caup.*IR (n=20, n=0.3467) clones. n.s. (not significant); Kruskal-Wallis test followed by Dunn’s many-to-one comparison test with Holm correction. (F and G) Wing discs bearing GFP-labeled Ras^V12^/*scrib^-/-^*+*lexA*.IR (F) and Ras^V12^/*scrib^-/-^*+*ara&mirr.*IR (G) clones stained with anti-β-galactosidase antibody for the *upd1-lacZ* reporter, dissected at 144-168 h AEL. (F”, F’’’, G”, and G’’’) Magnified images of the notum region of F, F’, G, and G’ respectively. (H) Quantification of the relative intensity of *upd1-lacZ* in Ras^V12^/*scrib^-/-^*+*lexA*.IR (n=11) and Ras^V12^/*scrib^-/-^*+*ara&mirr.*IR (n=8, p=0.0044) clones. **p < 0.01; Wilcoxon rank sum test. (I and J) Wing discs bearing GFP-labeled *lexA*.IR (I) and *ara&mirr.*IR (J) clones stained with anti-β-galactosidase antibody for the *upd1-lacZ* reporter, dissected at 120-144 h AEL. (I”, I’’’, J”, and J’’’) Magnified images of the notum region of I, I’, J, and J’ respectively. (K) Quantification of the relative intensity of *upd1-lacZ* in *lexA*.IR (n=16) and *ara&mirr.*IR (n=11, p=0.0004) clones. ***p < 0.001; Wilcoxon rank sum test. (L and M) Wing discs bearing GFP-labeled Ras^V12^/*scrib^-/-^*+*lexA*.IR (L), and Ras^V12^/*scrib^-/-^*+*ara&mirr.*IR (M) clones hybridized with *upd2* RNA probe to visualize *upd2* mRNA expression, dissected at 144-168 h AEL. (L”, L’’’, M”, and M’’’) Magnified images of the notum region of L, L’’, M, and M’ respectively. (N) Quantification of the relative intensity of *upd2* FISH signal in Ras^V12^/*scrib^-/-^*+*lexA*.IR (n=16) and Ras^V12^/*scrib^-/-^*+*ara&mirr.*IR (n=19, p=0.0026) clones. **p < 0.01; Wilcoxon rank sum test. (O and P) Wing discs bearing GFP-labeled Ras^V12^/*scrib^-/-^*+*lexA*.IR (O), and Ras^V12^/*scrib^-/-^*+*ara&mirr.*IR (P) clones hybridized with *upd3* RNA probe to visualize *upd3* mRNA expression, dissected at 144-168 h AEL. (O”, O’’’, P”, and P’’’) Magnified images of the notum region of O, O’, P, and P’ respectively. (Q) Quantification of the relative intensity of *upd3* FISH signal in Ras^V12^/*scrib^-/-^*+*lexA*.IR (n=13) and Ras^V12^/*scrib^-/-^*+*ara&mirr.*IR (n=21, p=0.0008) clones. ***p < 0.001; Wilcoxon rank sum test. Scale bar, 100 µm.

Importantly, the expression of Iro-C in the notum region is not uniform, rather their expression is enriched to the lateral notum (Figure S2F-S2H). We confirmed that the anti-cancer effect is indeed stronger in the Iro-C-positive lateral notum as the clone size of Ras^V12^*/scrib^-/-^* in the Caup-positive lateral notum was smaller than in the Caup-negative medial notum (Figure S2J, compare the lateral (left) and medial (right) areas, quantified in S2K), whereas wild-type clones showed no such difference between the two regions (Figure S2L, quantified in S2M). In addition, Mirr is intrinsically expressed in a specific portion of the hinge region (Figure S2F, magenta arrowheads), where the tumor growth of Ras^V12^*/scrib^-/-^*also appears to be suppressed (Figure S2I, magenta dotted lines). Together, these findings further support our conclusion that Iro-C-expressing fields possess an intrinsic anti-cancer property against IL-6-dependent malignant tumorigenesis.

Furthermore, Iro-C is expressed in a portion of the dorsal compartment in eye disc (Figure S3A-S3C), which is consistent with a previous report (McNeill et al, 1997). We found that clone size of Ras^V12^/*lgl*^-/-^ tumors was smaller in the *ara-lacZ*-positive area than in the *ara-lacZ*-negative area (Fig S3D, quantified in S3E) whereas wild-type clones were induced uniformly across both regions (Fig S3F, quantified in S3G). These findings suggest that the anti-cancerized field established by Iro-C is not restricted to the wing disc but may represent a conserved mechanism operating across *Drosophila* epithelial tissues.

Given that Iro-C acts as a transcriptional repressor, we hypothesized that Iro-C represses genes required for malignant tumor growth, such as the cytokine *unpaired* (*upd1*), a *Drosophila* homolog of interleukin-6 (IL-6) (Harrison *et al*, 1998; Mukherjee *et al*, 2005; Wu *et al*, 2010). As reported previously (Doggett *et al*, 2015; Külshammer *et al*, 2015), *upd1* was upregulated in Ras^V12^*/scrib^-/-^* clones in the P+H region at both the transcriptional and protein levels (Figure 3F and S4A). In contrast, Upd1 expression was markedly suppressed in Ras^V12^*/scrib^-/-^* cells within the notum (Figure 3F and S4A). Strikingly, knockdown of the Iro-C genes *ara* and *mirr* in Ras^V12^*/scrib^-/-^*cells upregulated Upd1 expression in the notum (Figure 3G, quantified in 3H, Figure S4B, quantified in S4C). Moreover, knockdown of *ara* and *mirr* in wild-type cells also resulted in the upregulation of *upd1* in the notum region (Figure 3I and 3J, quantified in 3K), as well as in the part of hinge region where *mirr* is normally expressed (Figure 3J’, yellow arrowheads and S2F, magenta arrowheads), while it did not affect other tumor-promoting pathways such as Hippo or Wg (Figure S5A, S5B, S5D, and S5E, quantified in S5C and S5F). Furthermore, *upd2* and *upd3* were also upregulated by the knockdown of *ara* and *mirr* in Ras^V12^*/scrib^-/-^* clones in the notum (Figure 3L, 3M, 3O, 3P, S3D, and S4E, quantified in 3N, 3Q and S4F respectively). As Iro-C proteins primarily act as transcriptional repressors (Bilioni *et al*, 2005; Carrasco-Rando *et al*, 2011), they could intrinsically repress the transcription of *upd1, upd2* and *upd3*. Indeed, we identified multiple ACAnnTGT motifs, the small palindromic sequence defined as the minimal DNA binding site of Iro-C (Bilioni *et al*., 2005), within the putative regulatory regions of the *upd1, upd2,* and *upd3* genes (Figure S4G), suggesting that Iro-C represses the transcription of these genes. These findings suggest that the notum-specific expression of Iro-C represses *upds* transcription, thereby creating anti-cancerized field against malignant tumorigenesis.

### Impaired JAK-STAT activation by Iro-C creates anti-cancerized field in the notum

Upd proteins commonly activate the Janus kinase/signal transducer and activator of transcription (JAK-STAT) signaling pathway via the Domeless receptor (Dome), analogous to mammalian IL-6 receptor (Brown *et al*, 2001). Loss of *upd1*, *upd2, upd3*, *dome*, or *stat92E* (a STAT3 homolog (Hou *et al*, 1996; Yan *et al*, 1996)) significantly suppressed Ras^V12^*/scrib^-/-^* tumor growth in the P+H regions (Figure 4A-4H, quantified in 4I, S6A and S6B, quantified in S6C). On the other hand, knockdown or overexpression of these genes in the wild-type background did not affect cell proliferation (Figure S6D-S6K, quantified in S6L, Figure S6M-S6O, quantified in S6P). These data indicate that Ras^V12^*/scrib^-/-^* cells in the P+H regions activate JAK/STAT signaling through Upds upregulation, which is specifically required for tumor growth.

**Figure 4.**
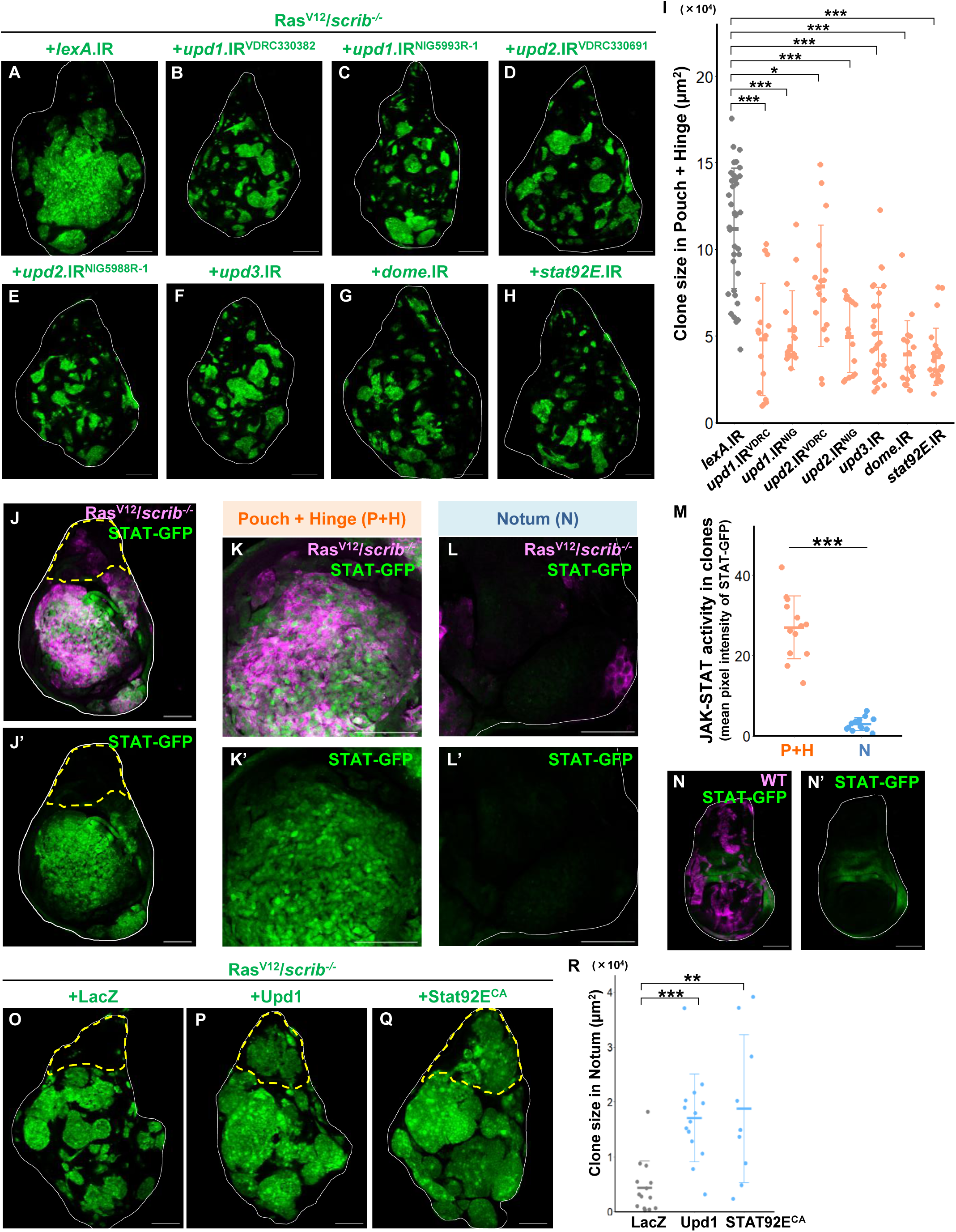
Impaired JAK-STAT activation by Iro-C creates anti-cancerized field in the notum. (A-F) Wing discs bearing GFP-labeled Ras^V12^/*scrib^-/-^*+*lexA*.IR (A), Ras^V12^/*scrib^-/-^*+*upd1*.IR^VDRC330382^ (B), Ras^V12^/*scrib^-/-^*+*upd1.*IR^NIG5993R-1^ (C), Ras^V12^/*scrib^-/-^*+*upd2.*IR^VDRC330691^ (D), Ras^V12^/*scrib^-/-^*+*upd2.*IR^NIG5988R-1^ (E), Ras^V12^/*scrib^-/-^*+*upd3.*IR (F), Ras^V12^/*scrib^-/-^*+*dome.*IR (G), and Ras^V12^/*scrib^-/-^*+*stat92E.*IR (H) clones, dissected at 120-144 h AEL. (I) Quantification of the clone size in the Pouch+Hinge (P+H) region for Ras^V12^/*scrib^-/-^*+*lexA*.IR (n=36), Ras^V12^/*scrib^-/-^*+*upd1*.IR^VDRC330382^ (n=16, p<0.0001), Ras^V12^/*scrib^-/-^*+ *upd1.*IR^NIG5993R-1^ (n=17, p<0.0001), Ras^V12^/*scrib^-/-^*+ *upd2.*IR^VDRC330691^ (n=17, p=0.0353), Ras^V12^/*scrib^-/-^*+ *upd2.*IR^NIG5988R-1^ (n=15, p<0.0001), Ras^V12^/*scrib^-/-^*+*upd3.*IR (n=27, p<0.0001), Ras^V12^/*scrib^-/-^*+*dome.*IR (n=17, p<0.0001), and Ras^V12^/*scrib^-/-^*+*stat92E.*IR (n=23, p<0.0001) clones. ***p < 0.001; *p < 0.05; Kruskal-Wallis test followed by Dunn’s many-to-one comparison test with Holm correction. (J-L) STAT-GFP expression in a wing disc bearing RFP-labeled Ras^V12^/*scrib^-/-^* clones (J). Magnified images of the Pouch+Hinge (P+H) region (K) or Notum (N) region (L), dissected at 144-168 h AEL. (M) Quantification of the mean pixel intensity of STAT-GFP in Ras^V12^/*scrib^-/-^* clones (n=13, p<0.0001) in the P+H or N region. ***p < 0.001; Wilcoxon rank sum test. (N) STAT-GFP expression in a wing disc bearing RFP-labeled WT clones, dissected at 96-120 h AEL. (O-Q) Wing discs bearing GFP-labeled Ras^V12^/*scrib^-/-^*+LacZ (O), Ras^V12^/*scrib^-/-^*+Upd1 (P), and Ras^V12^/*scrib^-/-^*+Stat92E^CA^ (Q) expressing clones, dissected at 144-168 h AEL. (R) Quantification of the clone size in the notum region for Ras^V12^/*scrib^-/-^*+LacZ (n=14), Ras^V12^/*scrib^-/-^*+Upd1 (n=14, p=0.0005), and Ras^V12^/*scrib^-/-^*+Stat92E^CA^ (n=9, p=0.0021) expressing clones. ***p < 0.001; **p < 0.01; Kruskal-Wallis test followed by Dunn’s many-to-one comparison test with Holm correction. Scale bar, 100 µm.

We then examined JAK-STAT activity in Ras^V12^*/scrib^-/-^*clones using the STAT-GFP reporter, which was strongly upregulated in the P+H regions but was significantly suppressed within the notum (Figure 4J-4L, quantified in 4M). In wild-type wing discs, STAT-GFP expression was restricted to the hinge region (Figure 4N) (Bach *et al*, 2007). Similarly, Ras^V12^/*scrib*.IR clones induced by *pdm2*-Gal4, a Gal4 driver specific to the pouch and part of the hinge region, exhibited strong upregulation of STAT-GFP (Figure S7A and S7B, quantified in S7E), whereas they hardly upregulate STAT-GFP when induced by *mirr*-Gal4, which is specific to the lateral part of the notum region (Figure S7C and S7D, quantified in S7E). Consistent with these data, forced activation of JAK-STAT signaling by overexpression of Upd1 or constitutively activated form of Stat92E (Stat92E^CA^) in Ras^V12^*/scrib^-/-^* clones caused significant overgrowth even in the notum (Figure 4O-4Q, quantified in 4R). These results indicate that impaired JAK-STAT activation by Iro-C creates anti-cancerized field against malignant tumorigenesis in the notum.

### Forced expression of Iro-C suppresses tumorigenesis in the tumor-prone regions

Finally, we examined whether forced expression of Iro-C could suppress tumor growth in the ‘tumor-prone’ pouch and hinge regions. Notably, overexpression of Iro-C in Ras^V12^*/scrib^-/-^* clones significantly reduced the tumor size in the P+H regions (Figure 5A-5D, quantified in 5E), while it only mildly affected wild-type tissue growth (Figure S8A-S8D, quantified in S8E). Furthermore, overexpression of an Iro-C protein Caup in Ras^V12^*/scrib^-/-^* clones suppressed the expression of *upd1, upd2,* and *upd3,* as well as the tumor growth in the P+H regions (Figure 5G, 5K, 5O, S9B and S9G, quantified in 5I, 5M, 5Q, S9E, and S9I, respectively). In contrast, overexpression of homeodomain mutant Caup (Caup^HD*^) lacking DNA-binding activity did not suppress *upds* expressions and tumorigenesis (Figure 5H, 5L, 5P, S9C, and S9H, quantified in 5I, 5M, 5Q, S9D, S9E and S9I, respectively). Together, these results indicate that expression of Iro-C is necessary and sufficient for creating anti-cancerized field against Upd/IL-6-dependent malignant tumorigenesis (Figure 6).

**Figure 5.**
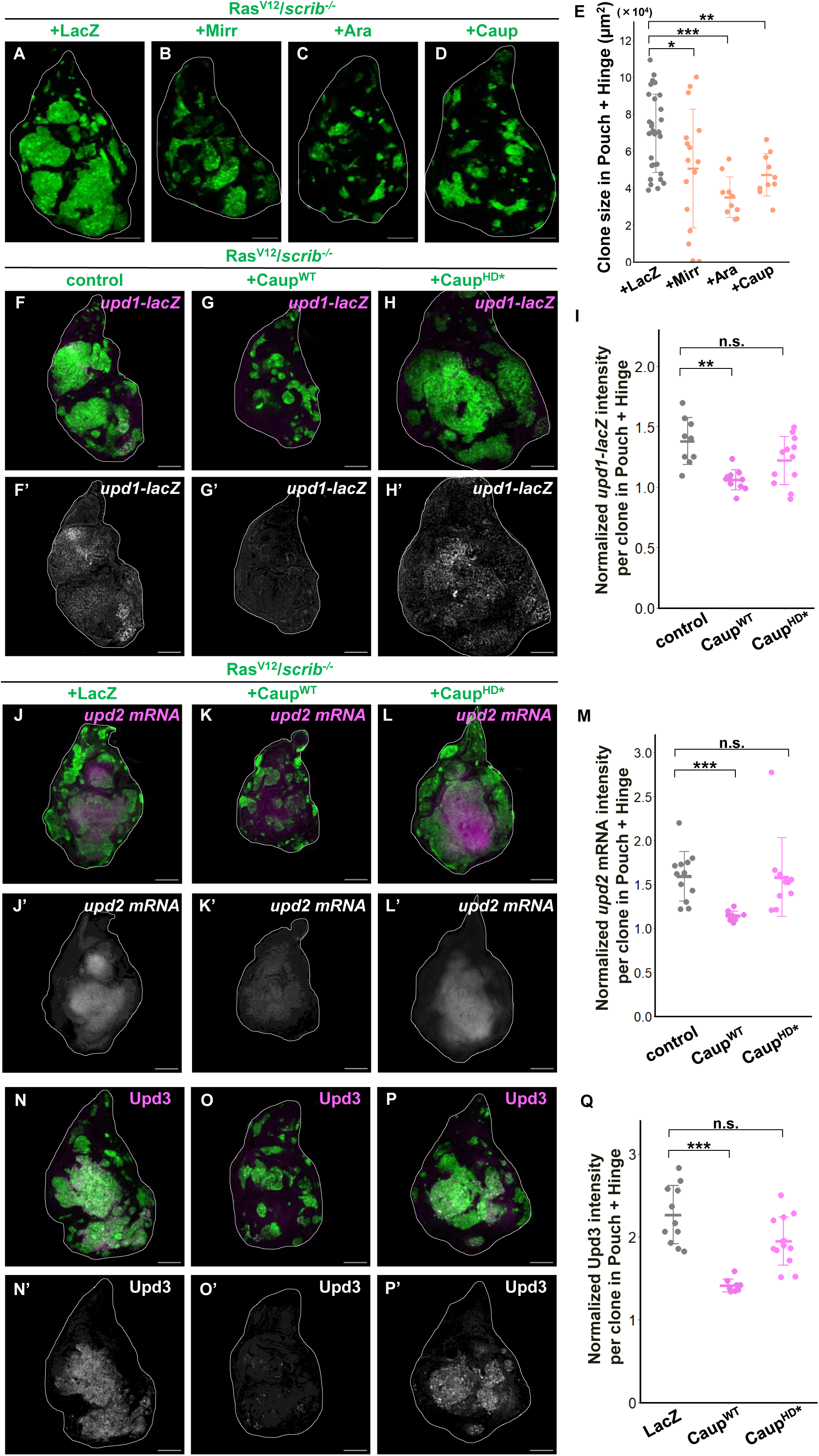
Forced expression of Iro-C suppresses tumorigenesis in the tumor-prone regions. (A-D) Wing discs bearing GFP-labeled Ras^V12^/*scrib^-/-^*+LacZ (A), Ras^V12^/*scrib^-/-^*+Mirr (B), Ras^V12^/*scrib^-/-^*+Ara (C), and Ras^V12^/*scrib^-/-^*+Caup (D) expressing clones, dissected at 120-144 h AEL. (E) Quantification of the clone size in the Pouch+Hinge (P+H) region for Ras^V12^/*scrib^-/-^*+LacZ (n=33), Ras^V12^/*scrib^-/-^*+Mirr (n=16, p=0.0121), Ras^V12^/*scrib^-/-^*+Ara (n=10, p<0.0001), and Ras^V12^/*scrib^-/-^*+Caup^WT^ (n=10, p=0.0095) expressing clones. ***p < 0.001; **p < 0.01; *p < 0.05; Kruskal-Wallis test followed by Dunn’s many-to-one comparison test with Holm correction. (F-H) Wing discs bearing GFP-labeled Ras^V12^/*scrib^-/-^* (F), Ras^V12^/*scrib^-/-^*+Caup^WT^ (G), and Ras^V12^/*scrib^-/-^*+Caup^HD*^ (H) expressing clones stained with anti-β-galactosidase antibody for the *upd1-lacZ* reporter, dissected at 144-168 h AEL. (I) Quantification of the relative intensity of *upd1-lacZ* for Ras^V12^/*scrib^-/-^*(n=9), Ras^V12^/*scrib^-/-^*+Caup^WT^ (n=11, p=0.0012), and Ras^V12^/*scrib^-/-^*+Caup^HD*^ (n=12, p=0.1396) expressing clones. **p < 0.01; n.s. (not significant); Kruskal-Wallis test followed by Dunn’s many-to-one comparison test with Holm correction. (J-L) Wing discs bearing GFP-labeled Ras^V12^/*scrib^-/-^*+LacZ (J), Ras^V12^/*scrib^-/-^*+Caup^WT^ (K), and Ras^V12^/*scrib^-/-^*+Caup^HD*^ (L) expressing clones hybridized with *upd2* RNA probe to visualize *upd2 mRNA* expression, dissected at 144-168 h AEL. (M) Quantification of the relative intensity of *upd2* FISH signal in Ras^V12^/*scrib^-/-^*+LacZ (n=12), Ras^V12^/*scrib^-/-^*+Caup^WT^ (n=10, p<0.0001), and Ras^V12^/*scrib^-/-^*+Caup^HD*^ (n=10, p=0.4883) expressing clones. ***p < 0.001; n.s. (not significant); Kruskal-Wallis test followed by Dunn’s many-to-one comparison test with Holm correction. (N-P) Wing discs bearing GFP-labeled Ras^V12^/*scrib^-/-^*+LacZ (N), Ras^V12^/*scrib^-/-^*+Caup^WT^ (O), and Ras^V12^/*scrib^-/-^*+Caup^HD*^ (P) expressing clones stained with anti-Upd3 antibody, dissected at 144-168 h AEL. (Q) Quantification of the relative intensity of Upd3 antibody for Ras^V12^/*scrib^-/-^*+LacZ (n=11), Ras^V12^/*scrib^-/-^*+Caup^WT^ (n=9, p<0.0001), and Ras^V12^/*scrib^-/-^*+Caup^HD*^ (n=13, p=0.11448) expressing clones. ***p < 0.001; n.s. (not significant); Kruskal-Wallis test followed by Dunn’s many-to-one comparison test with Holm correction.

**Figure 6.**
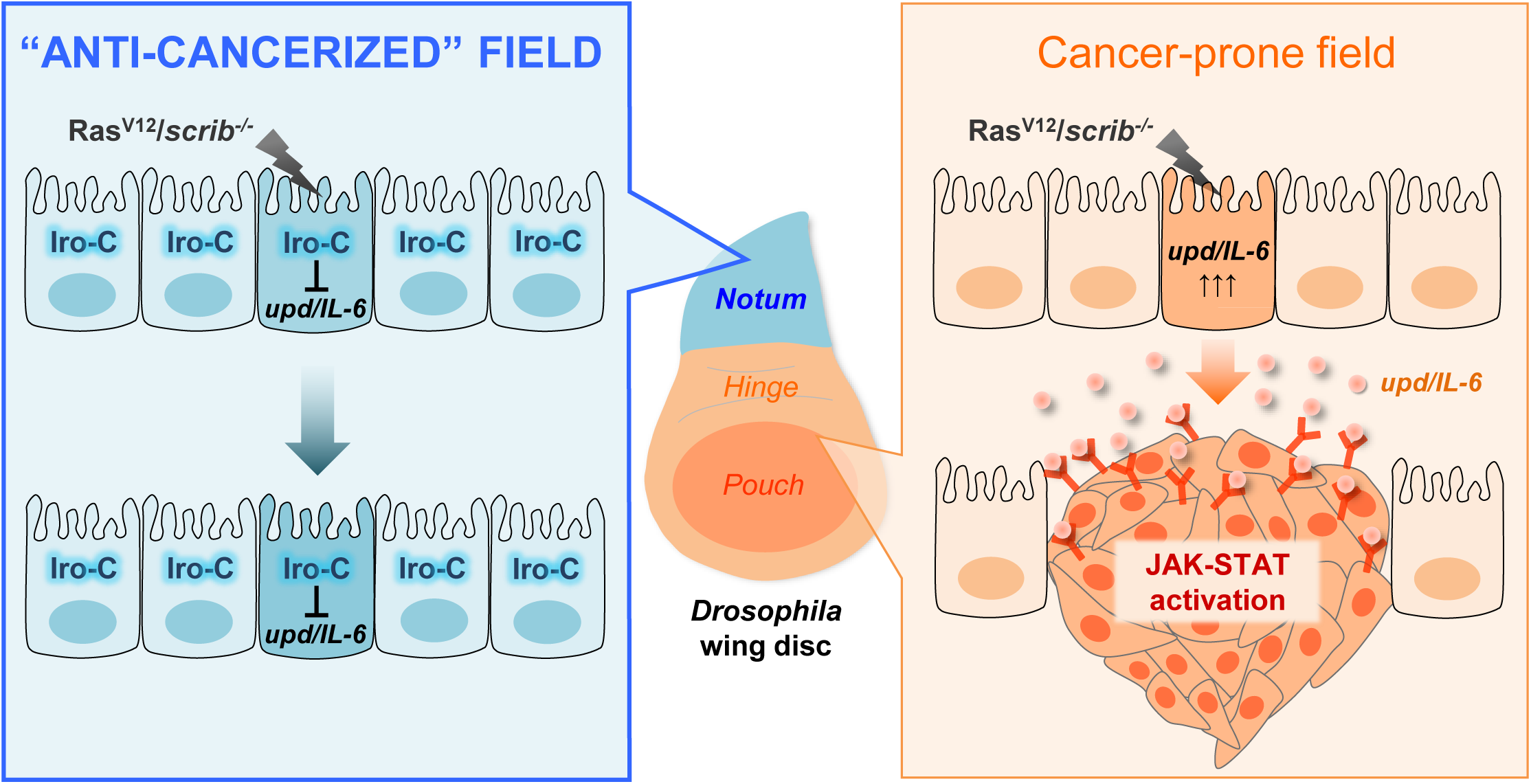
A model for Iro-C/IRX-mediated formation of anti-cancerized field against Upd/IL-6-dependent tumorigenesis. In the tumor-prone pouch and hinge regions, Ras^V12^/*scrib^-/-^*cells activate JAK/STAT signaling through upregulation of Upd1, Upd2 and Upd3, leading to the tumor formation. In contrast, in the anti-cancerized notum region, where Iro-C is intrinsically expressed, Iro-C represses the transcription of *upds* in Ras^V12^/*scrib^-/-^*cells, thereby preventing JAK/STAT activation and tumorigenesis.

## Discussion

Our genetic study in *Drosophila* identified Iro-C as a novel type of tumor suppressor that creates an intrinsic anti-cancerized field against Upd/IL-6-dependent malignant tumorigenesis. We found that Iro-C specifically suppresses growth of malignant tumors caused by Ras activation and cell polarity defect, but not benign tumors caused by Ras activation, YAP/Yki activation, or Tor activation. Thus, Iro-C, an evolutionarily conserved determinant of cell specification during normal development, also acts as a fail-safe mechanism against lethal, malignant tumorigenesis. Notably, forced expression of Iro-C significantly reduced the size of Ras^V12^*/scrib^-/-^* tumors in the tumor-prone regions. Given that cancers hijack developmental programs to progress toward malignancy, our observations suggest that manipulations of developmental cell-fate determinants could provide a novel, effective anti-cancer strategy.

Based on our previous and current data, JNK exerts a pro-growth effect on Ras^V12^/*scrib*^-/-^ tumors when JAK-STAT signaling is activated, as inhibition of JNK suppresses tumor growth (Igaki *et al*., 2006; Uhlirova & Bohmann, 2006). In contrast, in the notum region, where Upd-JAK-STAT signaling is intrinsically blocked by Iro-C, elevated JNK signaling exerts an anti-growth effect on Ras^V12^/*scrib*^-/-^ tumors (Figure S10). These findings suggest that the pro-growth versus anti-growth function of JNK in Ras^V12^/*scrib*^-/-^ tumors depends on the status of JAK-STAT signaling. Thus, JAK-STAT signaling switches the anti-growth effect of JNK to pro-growth effect, thereby enabling Ras^V12^/*scrib*^-/-^ tumor overgrowth and rendering these tumors highly dependent on—or “addicted” to—JAK-STAT signaling. In contrast, hyperplastic Ras^V12^ tumors exhibit only low levels of JAK-STAT and JNK activity (Enomoto *et al*, 2021; Igaki *et al*., 2006), and are therefore “non-addicted” to JAK-STAT signaling, which could explain why hyperplastic tumors can develop in the notum region.

Upd/IL-6-JAK-STAT signaling has been shown to be essential for tissue regeneration in *Drosophila* (Jiang *et al*, 2009) and vertebrates (Cressman *et al*, 1996). In *Drosophila* wing disc, genetic ablation of the pouch cells induces regenerative response to surrounding living cells, which cause Upd1- and Upd3-mediated JAK-STAT activation (Santabárbara-Ruiz *et al*, 2015), whereas genetic ablation of the notum cells does not induce regeneration (Martín *et al*, 2017). These findings suggest that Iro-C-mediated repression of *upds* in the notum may restrict not only its tumorigenic potential but also its regenerative capacity. Notably, a recent work in human pancreatic cancer has shown that highly regenerative cell populations preferentially tend to undergo clonal expansion and neoplastic transformation upon acquiring oncogenic mutations (Neuhöfer *et al*, 2021). Thus, the intrinsic sensitivity of epithelial cells to Upd/IL-6-JAK-STAT activation provides an explanation for how they acquire both tumorigenic potential and regenerative capacity.

The IL-6-STAT3 signaling pathway plays an important role in the progression of multiple human cancers, including gastric, breast, liver, colorectal, ovarian, pancreatic, lung, and kidney cancers (Huang *et al*, 2022; Kamińska *et al*, 2015). In renal cell carcinoma (RCC), which originates from the renal tubular epithelium of the kidney, IL-6-STAT3 activation occurs in an autocrine manner and promotes tumorigenesis (Kamińska *et al*., 2015). Notably, most RCC tumors arise from proximal tubule cells or intercalated cells of the collecting duct, but are rarely derived from distal nephron segments, including loop of Henle, distal convoluted tubule, and macula densa (Lindgren *et al*., 2017). Intriguingly, IRX1, a vertebrate homolog of Iro-C, is specifically expressed in these areas with low tumor incidence (Lindgren *et al*., 2017; Lindström *et al*, 2021). Thus, IRX1 may create an intrinsic anti-cancerized field against IL-6-dependent RCC tumors in these nephron segments. It has also been reported that overexpression of IRX1 suppresses proliferation of human cholangiocarcinoma cells (Jung *et al*, 2019) and human gastric cancer cells (Guo *et al*, 2010). Together, our study indicates that Iro-C/IRX family proteins act as evolutionary conserved tumor suppressors and could serve as a potential target for cancer prevention and therapy against IL-6-dependent tumorigenesis by creating an anti-cancerized field.

## Material & Methods

### Drosophila strains

*Drosophila melanogaster* strains were raised in vials containing a standard cornmeal-sucrose-yeast food, maintained at 25&#x25A1; xsunless otherwise stated. The sex of larvae dissected for most imaginal disc studies was not differentiated. For producing fluorescently labeled mitotic clones in larval imaginal discs, the following tester strains were used for MARCM analysis; “*ubxFLP; Act>y^+^>Gal4, UAS-GFP; FRT82B, tublin-Gal80*”, “*upd1-lacZ; ubxFLP, Act>y^+^>Gal4, UAS-GFP; FRT82B, tublin-Gal80*”, “*ubxFLP; tublin-Gal80, FRT40A, UAS-Ras^V12^; Act>y^+^>Gal4, UAS-GFP*”, and “*ubxFLP; tublin-Gal80, FRT40A, UAS-Ras^V12^; Act>y^+^>Gal4, UAS-GFP, UAS-scrib.IR*”. The following fly strains were used for each experiment: *STAT-GFP* (Bach et al., 2007), *UAS-Ras^V12^* (Igaki et al., 2006), *UAS-Upd1* (E. Bach), *UAS-3HA-Stat92E^ΔNΔC^* (Ekas et al, 2010), *upd1-lacZ* (Chao et al, 2004), *ara^rF209^* (*ara-lacZ*)*, UAS-Ara* (Gomez-Skarmeta et al, 1996), *UAS-Caup^WT^* (Barrios et al, 2015), *UAS-Caup^HD*^* (Barrios et al., 2015), *UAS-Mirr* (McNeill *et al*., 1997), *scrib^1^* (Bilder et al, 2000), *dlg^m52^* (Goode & Perrimon, 1997), *lgl^4^* (Gateff & Schneiderman, 1967), *UAS-CD8-PARP-Venus* (Williams et al, 2006), *UAS-miRHG* (Siegrist et al, 2010), *UAS-Bsk^DN^* (Adachi-Yamada *et al*, 1999); *UAS-EGFR^9534^, UAS-Dcr2^24650^, UAS-lacZ^8529^, UAS-lexA-RNAi^67947^, UAS-ara&mirr-RNAi^42960^*, *UAS-dome-RNA^53890^*, *UAS-scrib-RNAi^35748^*, *Caup-GFP^56155^*, *Mirr-GFP^68183^*, *upd3^Delta^ ^55728^*, *Tsc1^Q600X^ ^82163^*, *GMR76A01-Gal4^46953^*, *mirr^DE^-Gal4^29650^, Diap1^j5C8^ (diap1-lacZ)^12093^* (Bloomington Drosophila Stock Center); *UAS-caup-RNAi^2933^*, *UAS-upd1-RNAi^330382^*, *UAS-upd2-RNAi^330691^*, *UAS-upd3-RNAi^27134^* (Vienna Drosophila Resource Center); *UAS-stat92E-RNAi^4257R-2^, UAS-upd1-RNAi^5993R-1^, UAS-upd2-RNAi^5988R-1^* (National Institute of Genetics).

### Immunofluorescence and confocal imaging

Wandering third instar larvae (the exact timing of dissection for each experiment is described in the Figure Legends) were dissected and fixed with 4% paraformaldehyde (PFA) in PBS (nacalai tesque; 09154-85) for 20 min at room temperature and washed with PBS containing 0.1% Triton X-100 (nacalai tesque; 35501-15) solution (PBT) three times. Following PBT washes, samples were blocked with PBT containing 5% normal donkey serum (nacalai tesque; 017-000-121) (PBTn) for 30 min and then incubated with primary antibodies at 4°C overnight. Samples were incubated with Alexa Fluor-conjugated secondary antibodies (Thermo Fisher Scientific; 1:250) for 2 hours at room temperature, followed by washing with PBT three times. After PBT washes, samples were mounted with DAPI (Sigma-Aldrich; MBD0015)-containing mounting medium. Images were taken with a Leica STELLARIS 5 or TCS SP8 confocal microscope.

### Antibodies used

Larval tissues were stained with chicken anti-β-galactosidase antibody (Abcam; ab9361, 1:2000), chicken anti-GFP antibody (AVES Labs; GFP-1010, 1:1000), mouse anti-Wingless antibody (Developmental Studies Hybridoma Bank, 4D4, 1:100), rabbit anti-PH3 antibody (Cell Signaling Technology; 9701, 1:100), rabbit anti-cleaved-PARP antibody (Cell Signaling Technology; 9541, 1:100), rabbit anti-mCherry antibody (Abcam; ab167453, 1:250), rabbit anti-Upd1 antibody (D. Harrison, 1:500), rat anti-Upd3 antibody (Hirooka *et al*, 2025; Li *et al*, 2025). Secondary antibodies used are as follows: Goat anti-Chicken IgY (H+L) Cross-Adsorbed Secondary Antibody Alexa Fluor™ Plus 488, Goat anti-Chicken IgY (H+L) Cross-Adsorbed Secondary Antibody Alexa Fluor™ Plus 555, Alexa Fluor™ 546 goat anti-rabbit IgG (H+G), Goat anti-Rabbit IgG (H+L) Highly Cross-Adsorbed Secondary Antibody Alexa Fluor™ 647, Goat anti-Rat IgG (H+L) Antibody Alexa Fluor™ 647 Conjugated (Thermo Fisher Scientific; 1:250). RFP-Booster_ATTO594 (proteintech; rba594-100, 1:50) was used to enhance the RFP signal.

### Temperature-shift experiments

For experiments using the temperature-sensitive Gal80 (Gal80^ts^) system, larvae were reared at 18°C to maintain Gal4 suppression and then shifted to 29°C to induce UAS-driven transgene expression in the wing imaginal discs. For Figure S1, larvae were maintained at 18°C for 8 days and subsequently shifted to 29°C for 24 hours before dissection. For Figure S5A and S5C, larvae were maintained at 18°C for 6 days and shifted to 29°C for 48 hours before dissection. For Figure S5B and S5D, larvae were maintained at 18°C for 6 days and shifted to 29°C for 72 hours before dissection.

### Heat-shock-induced MARCM system

To induce cell clones using the heat-shock-induced MARCM system, flies were allowed to lay eggs for 8 hours. Parental flies were then removed, and larval progeny were heat-shocked at 37°C for 40 min at 48 h AEL to induce clones. Larvae were subsequently dissected at 89-97 h, 113-121 h, or 137-145 h AEL for analysis. GFP-labeled wild-type or Ras^V12^/*lgl^-/-^* clones were generated using the following tester line: *y, w, hsFLP; tub-Gal80, FRT40A; tub-Gal4, UAS-CD8GFP*.

### EdU incorporation

EdU incorporation was assayed using Click-iT Plus EdU Cell Proliferation Kit for Imaging, Alexa Fluor 647 dye (Thermo Fisher Scientific; c10640) following the manufacturer’s instructions: After larvae were dissected in Schneider’s *Drosophila* medium (Gibco; 21720024), EdU (10 µM final concentration) was incorporated into discs for 3 hours. Afterward, discs were fixed for 30 min in 4% PFA, then incubated in EdU click-iT reaction cocktail for 30 min.

### Fluorescence in situ hybridization (FISH)

Antisense RNA probes for *upd2* and *upd3* were synthesized by *in vitro* transcription using a DIG RNA labeling kit (Roche; 11175025910) according to the manufacturer’s instructions. The template DNA was amplified from *Drosophila melanogaster* genomic DNA, and purified RNA probes were cleaned using the NucleoSpin® RNA Clean-up XS kit (Macherey-Nagel) and eluted in RNase-free water. The concentration of RNA was adjusted to 0.1 µg/µL and stored at −80°C until use. Sense probes were synthesized in the same manner and used as negative controls.

Larvae were dissected in PBS, fixed with 4% PFA in PBS for 25 min at RT and 5 min on ice, and reacted with methanol for 5 min at −20°C. After rinsing twice with ethanol, the imaginal discs were permeated in a mixture of xylene and ethanol (1:1) for 1 hour on ice, hydrated with an ethanol series (100, 80, 50, and 25% ethanol) and RNase free water (nacalai tesque; 06442-95) on ice. After washing twice with ethanol, samples were permeabilized in 80% acetone for 10 min at −20°C, washed twice with PBS/0.1% Tween 20, and refixed with 4% PFA in PBS for 20 min at RT. After washing twice with PBS/0.1% Tween 20, samples were prehybridized in hybridization buffer [50% Formyl amide, 5× standard saline citrate (SSC), 100 µg /mL Salmon sperm DNA, 50 µg/mL heparin, 0.1% Tween 20] for 20 min at 50°C and hybridized in hybridization buffer containing 1.0 ng/µL DIG-labeled RNA probe for *upd2* or *upd3* for 14 hours at 50 °C. After washing twice with 2× SSC, samples were treated with Rnase A (MACHEREY-NAGEL; 740505.50) in Rnase buffer (20 μg/ml) [10 mM trisHCl (pH 8.0), 5 mM EDTA, 500 mM NaCl, and 0.1% Tween 20] for 10 min at RT and washed with SSC series (2× and 1× SSC) and PBT. After blocking with PBTn for 30 min at RT, samples were incubated overnight at 4°C with Chicken anti-GFP antibody (AVES Labs; GFP-1010, 1:1000). After washing three times with PBT, samples were again blocked with PBTn and incubated overnight at 4°C with Goat anti-Chicken IgY (H+L) Cross-Adsorbed Secondary Antibody, Alexa Fluor™ Plus 488 (Thermo Fisher Scientific; A-32931, 1:250). After washing three times with PBT, samples were blocked with 1×Blocking solution (Roche; 11096176001) and incubated for two hours at RT with sheep anti-Digoxigenin-AP Fab fragments (Roche; 11093274910, 1:2000). After washing twice with TBS-T (TBS and 0.1% Triton X-100), samples were incubated with BCIP NBT (nacalai tesque; 19880-84) for 4 hours at RT. Samples were mounted with DAPI-containing mounting medium. The reaction product generated by BCIP/NBT has fluorescence (Trinh *et al*, 2007), which were excited with a 633-nm laser and detected with a 650-nm long pass filter on a Leica STELLARIS 5 confocal microscope.

### Quantification and statistical analysis

#### Clone size

Clone size was quantified as either the total clone area or the total clone area / region area (notum or pouch + hinge) using Fiji software (Schindelin *et al*, 2012). The boundary between the notum and hinge regions was determined based on the proximal fold structure of the dorsal hinge, visualized by DAPI staining.

#### Signal intensity

Mean signal intensity within GFP-positive cells was measured as the mean pixel intensity of antibody staining, in situ hybridization signals, or reporter signals using Fiji software. The values were normalized to the mean pixel intensity of GFP-negative cells. For quantification of the STAT-GFP reporter, the intensity in RFP-positive cells was not normalized to that in RFP-negative cells, in order to avoid the influence of endogenous JAK-STAT activity in the hinge region.

#### Cell proliferation

The number of cells elevated PH3 or EdU signal in the GFP-positive clones was counted and normalized by DAPI staining in order to correct the cell number per unit area.

A series of quantitative data was plotted in RStudio (version 2025.05.1+513). For two-group comparisons, the Wilcoxon rank sum test was applied. For multiple comparisons, the Kruskal-Wallis test followed by Dunn’s many-to-one rank comparison test was used when comparing each group against a single control, or Dunn’s all-pairs rank comparison test when performing pairwise comparisons among all groups; p-values were adjusted using the Holm method. Significance of difference was represented by p-values (where n.s., non-significant difference; *p < 0.05, **p < 0.01, ***p < 0.001).

## Resource availability

### Lead contact

Further information and request for resources and reagents should be directed to and will be fulfilled by the lead contact, Tatsushi Igaki

### Materials availability

*Drosophila* lines generated in this study are available from the lead contact without restriction.

### Ethics declarations

The authors declare no competing interests.

### Data availability

All data reported in this paper will be shared by the lead contact upon request.

### Code availability

No custom code was used in this study.

### Author Contributions

T.N, M.E, and T.I. designed experiments; T.N conducted experiments with input from M.E. and T.I.; T.N, M.E, and T.I. analyzed the data; T.N, M.E, and T.I. wrote the manuscript; All authors reviewed the manuscript.

## Acknowledgements

The authors thank M. Matsuoka, M. Koijima and M. Sada for technical support; M. Matsuyama, T. Kobayashi, and members of the Igaki laboratory for discussions, D. Harrison for providing anti-Upd1 antibody; E. Bach, T. Xu, D. Bilder, H. Nakagoshi, S. Campuzano, the Bloomington Drosophila Stock Center (BDSC, Indiana, USA), the Vienna Drosophila Resource Center (VDRC, Vienna, Austria), the Drosophila Genomics and Genetic Resources (DGGR, Kyoto, Japan), and National Institute of Genetics (NIG, Shizuoka, Japan) for providing fly stocks. This work was supported in part by grants from the MEXT/JSPS KAKENHI (21H05284 and 21H05039 to T.I, 23K27366, 23K24080 and 23K18198 to M.E, 24KJ1515 to T.N), AMED-CREST, Japan Agency for Medical Research and Development (23gm1710002h0002) to T.I, Japan Science and Technology Agency. PRESTO (JPMJPR2141) to M.E, Princess Takamatsu Cancer Research Fund to M.E, The Foundation for Promotion of Cancer Research in Japan to M.E, and the Takeda Science Foundation to T.I, and M.E.

During the preparation of this work the author used ChatGPT 5.1 in order to improve the readability and language of the manuscript. After using this tool, the authors reviewed and edited the content as needed and take full responsibility for the content of the published article.

## Figure Legends

**Figure S1.**
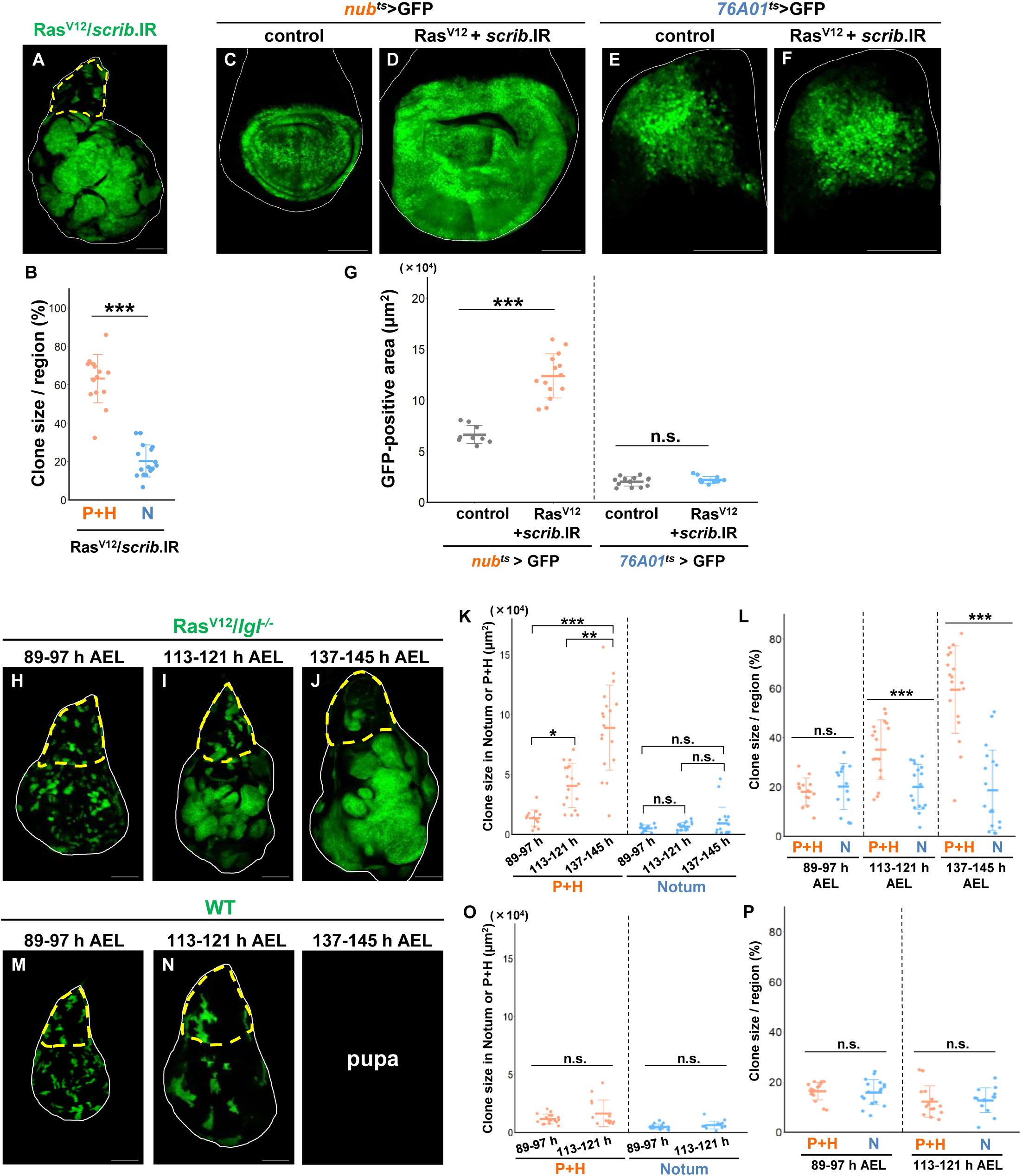
The notum region behaves as anti-cancerized field against malignant tumorigenesis, Related to Figure 1. (A) Wing discs bearing GFP-labeled Ras^V12^+*scrib*.IR*-*expressing clones dissected at 144-168 h AEL. (B) Quantification of the clone size in the Pouch+Hinge (P+H) or Notum (N) (% of the clone area per region area) for Ras^V12^+*scrib*.IR (n=16, p<0.0001) clones. ***p < 0.001; Wilcoxon rank sum test. (C-F) GFP-labeled control (C and E) and Ras^V12^+*scrib.*IR (D and F) cells induced by *nub*-Gal4 or *76A01*-Gal4. (G) Quantification of the GFP-positive area for *nub*^ts^>GFP (n=9), *nub*^ts^>GFP, Ras^V12^+*scrib.*IR (n=14, p<0.0001), *76A01*^ts^>GFP (n=13), and *76A01*^ts^>GFP, Ras^V12^+*scrib*.IR (n=12, p=0.6885). ***p < 0.001; n.s. (not significant); Wilcoxon rank sum test. (H-J) Wing disc bearing heat-shock-induced GFP-labeled Ras^V12^/*lgl^-/-^*clones, dissected at 89-97 h (H), 113-121 h (I), or 137-145 h (J) AEL. (D) Quantification of clone size in the P+H and notum regions at 89-97 h (n=14), 113-121 h (n=17), and 137-145 h (n=18) AEL. In the P+H region, clone size increased significantly over time (89-97 h vs. 113-121 h, p=0.0115; 113-121 h vs. 137-145 h, p=0.0069; 89-97 h vs. 137-145 h, p< 0.0001). In contrast, clone size in the notum region did not differ significantly across the three time points (89-97 h vs. 113-121 h, p=1; 113-121 h vs. 137-145 h, p=1; 89-97 h vs. 137-145 h, p=1). ***p < 0.001; **p < 0.01; n.s. (not significant); Kruskal-Wallis test followed by Dunn’s all-pairs comparison test with Holm correction. (E) Quantification of the Ras^V12^/*lgl^-/-^*clone size per region area, dissected at 89-97 h (n=14, p=0.4213), 113-121 h (n=17, p=0.0007), or 137-145 h AEL (n=18, p<0.0001). ***p < 0.001; n.s. (not significant); Wilcoxon rank sum test. (F-H) Wing disc bearing heat-shock-induced GFP-labeled wild-type clones, dissected at 89-97 h (M) or 113-121 h (N). (I) Quantification of the wild-type clone size, dissected at 89-97 h (n=17) and 113-121 h (n=14, P+H: p=0.6480, N: p=0.1265) AEL. n.s. (not significant); Wilcoxon rank sum test. (J) Quantification of the wild-type clone size per region area, dissected at 89-97 h (n=17, p=0.3953) or 113-121 h AEL (n=14, p=0.5124). n.s. (not significant); Wilcoxon rank sum test. Scale bar, 100 µm.

**Figure S2.**
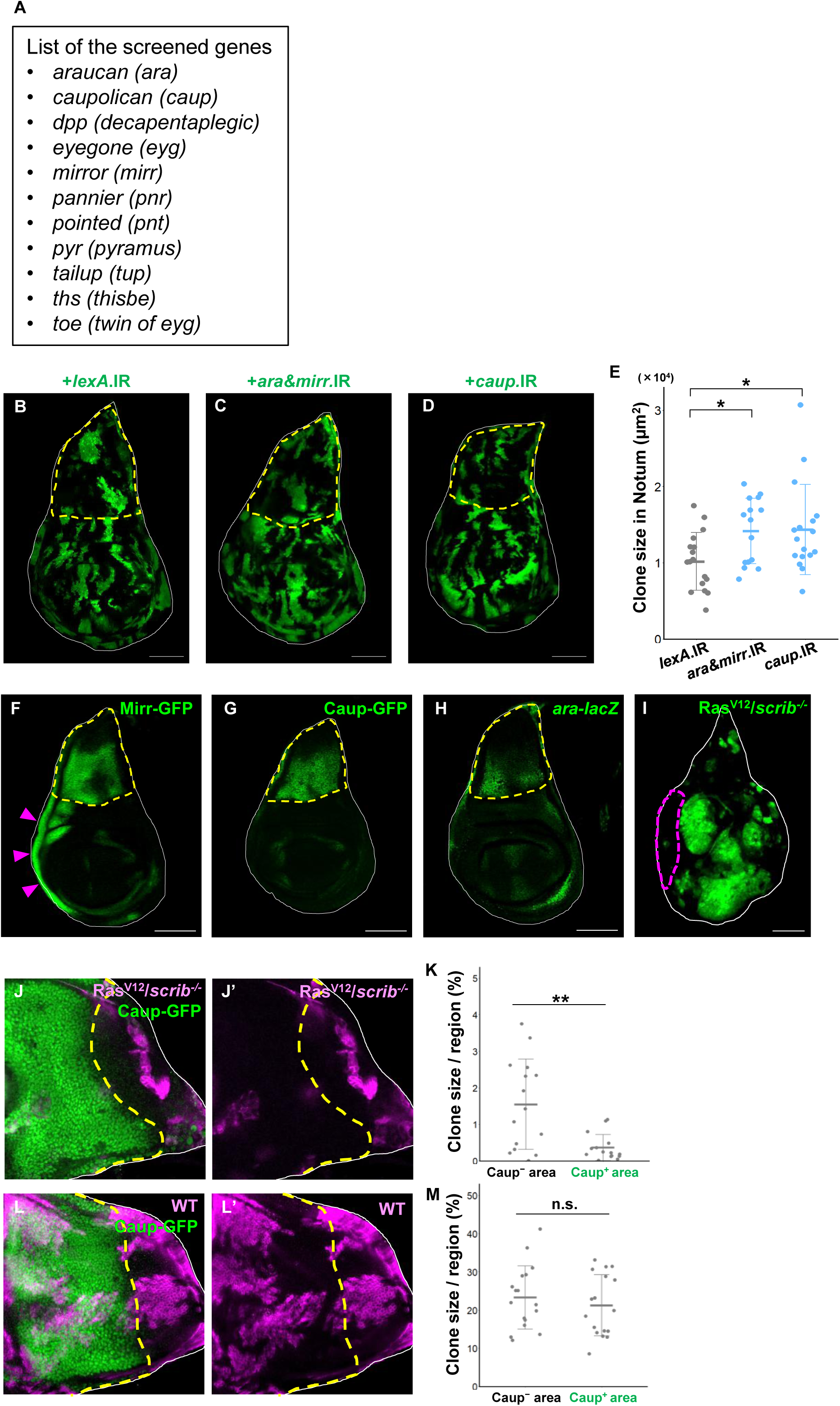
Iro-C is expressed in the notum region, Related to Figure 3. (A) A list of the screened genes that were knocked down in Ras^V12^/*scrib^-/-^* cells to identify the factors responsible for creating anti-cancerized field. (B-D) Wing discs bearing GFP-labeled *lexA*.IR (B), *ara&mirr.*IR (C), and *caup.*IR (D) clones, dissected at 96-120 h AEL. (E) Quantification of the clone size in the notum region for *lexA*.IR (n=17), *ara&mirr.*IR (n=15, p=0.0403), and *caup.*IR (n=17, p=0.0403) clones. *p < 0.05; Kruskal-Wallis test followed by Dunn’s many-to-one comparison test with Holm correction. (F-H) Expression patterns of Mirr-GFP (F), Caup-GFP (G), and *ara-lacZ* (H) in wing discs, dissected at 96-120 h AEL. (I) Wing disc bearing GFP-labeled Ras^V12^/*scrib^-/-^* cells, dissected at 144-168 h AEL. (J and L) Magnified images of the notum region of wing disc bearing RFP-labeled Ras^V12^/*scrib^-/-^*(J) or wild-type (L) clones, dissected at 120-144 h (J) or 96-120 h (L) or AEL. Caup-expressing area is labeled by Caup-GFP. (K and M) Quantification of the clone size of Ras^V12^/*scrib^-/-^* (K, n=15, p=0.0079) or wild-type (M, n=17, p=0.6297) clones in Caup-GFP-positive or -negative region (% of the clone area per region area). **p < 0.01; n.s. (not significant); Wilcoxon rank sum test. Scale bar, 100 µm.

**Figure S3.**
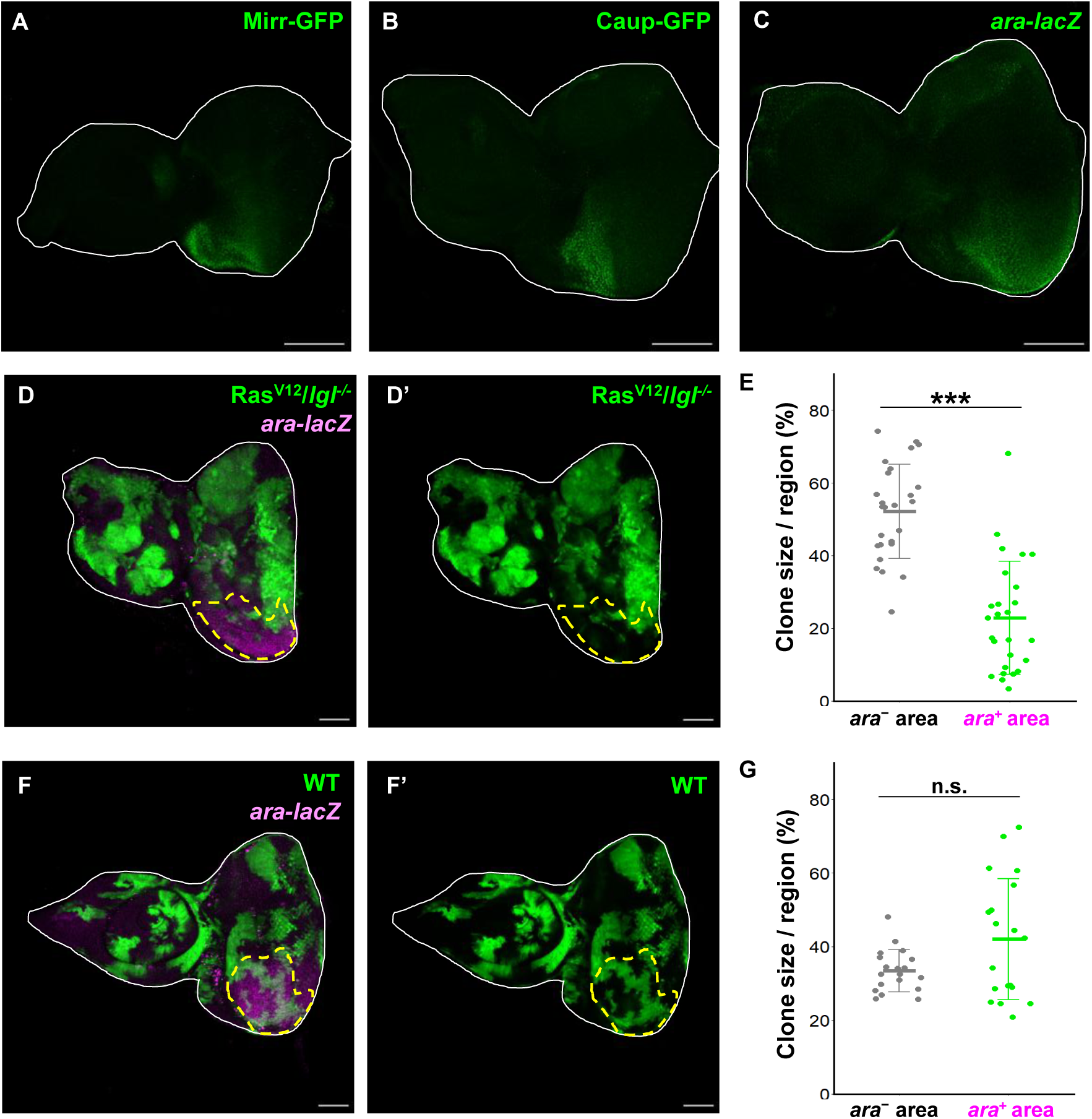
Iro-C also establishes anti-cancerized field against malignant tumors in the eye disc. (A-C) Expression patterns of Mirr-GFP (A), Caup-GFP (B), and *ara-lacZ* (C) in eye-antennal discs, dissected at 96-144 h AEL. (D and F) Eye-antennal discs bearing GFP-labeled Ras^V12^/*lgl*^-/-^ (D) or wild-type (F) clones. *ara*-expressing region is labeled by *ara-lacZ*. (E and G) Quantification of the clone size of Ras^V12^/*lgl*^-/-^ (E, n=26, p<0.0001) or wild-type (G, n=19, p=0.2801) or clones in *ara-lacZ*-positive or -negative region (% of the clone area per region area). ***p < 0.001; n.s. (not significant); Wilcoxon rank sum test. Scale bar, 100 µm.

**Figure S4.**
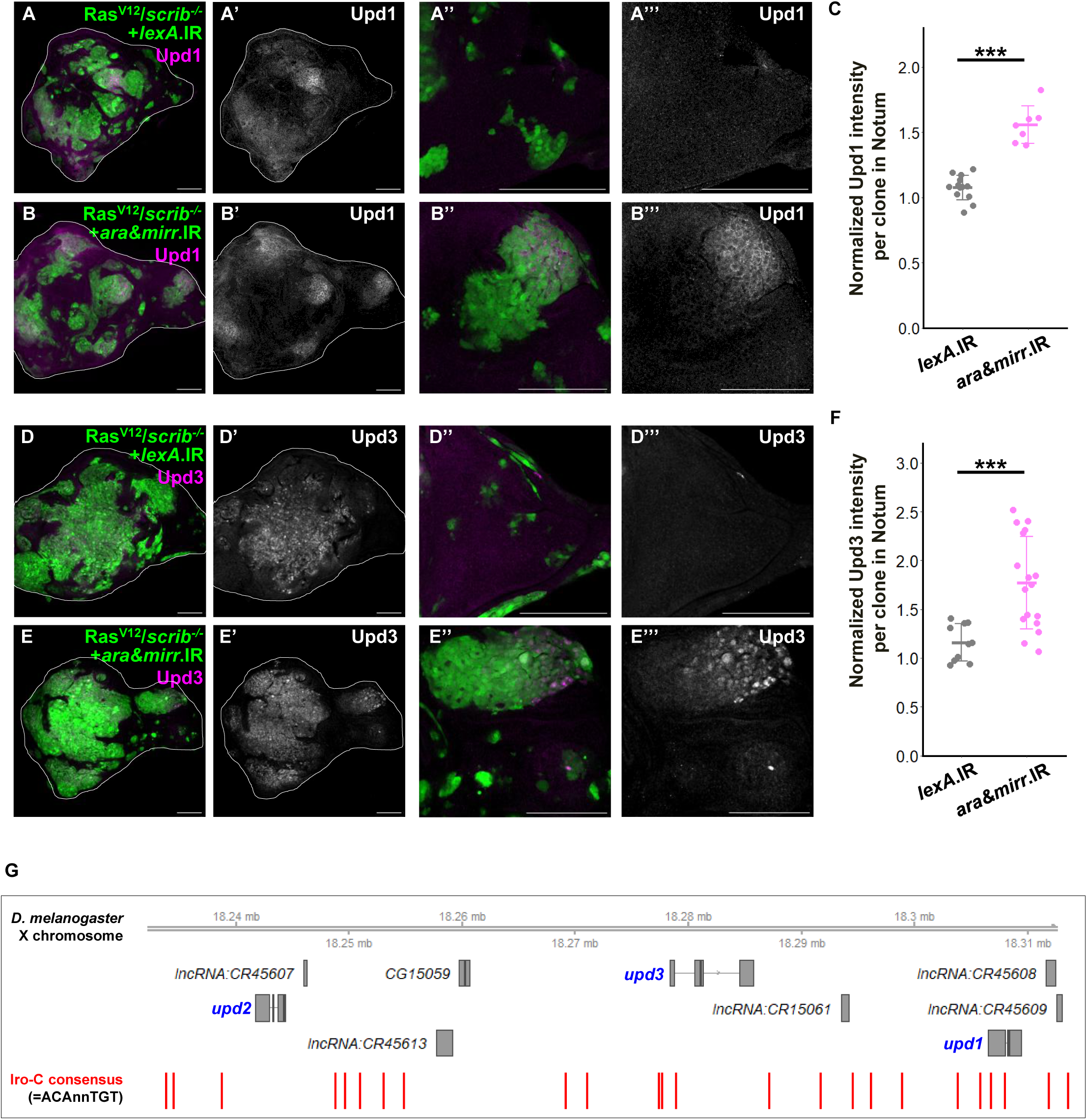
Notum-specific expression of Iro-C represses *upd/IL-6* transcription, Related to Figure 3. (A and B) Wing discs bearing GFP-labeled Ras^V12^/*scrib^-/-^*+*lexA*.IR (A), and Ras^V12^/*scrib^-/-^*+*ara&mirr.*IR (B) clones stained with anti-Upd1 antibody, dissected at dissected at 120-168 h AEL. (A”, A’’’, B”, and B’’’) Magnified images of the notum region of A, A’, B, and B’ respectively. (C) Quantification of the relative intensity of Upd1 antibody in Ras^V12^/*scrib^-/-^*+*lexA*.IR (n=13) and Ras^V12^/*scrib^-/-^*+*ara&mirr.*IR (n=7, p=0.0004) clones. ***p < 0.001; Wilcoxon rank sum test. (D and E) Wing discs bearing GFP-labeled Ras^V12^/*scrib^-/-^*+*lexA*.IR (D), and Ras^V12^/*scrib^-/-^*+*ara&mirr.*IR (E) clones stained with anti-Upd3 antibody, dissected at dissected at 144-168 h AEL. (D”, D’’’, E”, and E’’’) Magnified images of the notum region of D, D’, E, and E’ respectively. (F) Quantification of the relative intensity of Upd3 antibody in Ras^V12^/*scrib^-/-^*+*lexA*.IR (n=10) and Ras^V12^/*scrib^-/-^*+*ara&mirr.*IR (n=17, p=0.0004) clones. ***p < 0.001; Wilcoxon rank sum test. Scale bar, 100 µm. (G) Physical map of the *upd1, upd2,* and *upd3* locus. Iro-C consensus motifs (ACAnnTGT, red bars) are visualized using the R package Gviz.

**Figure S5.**
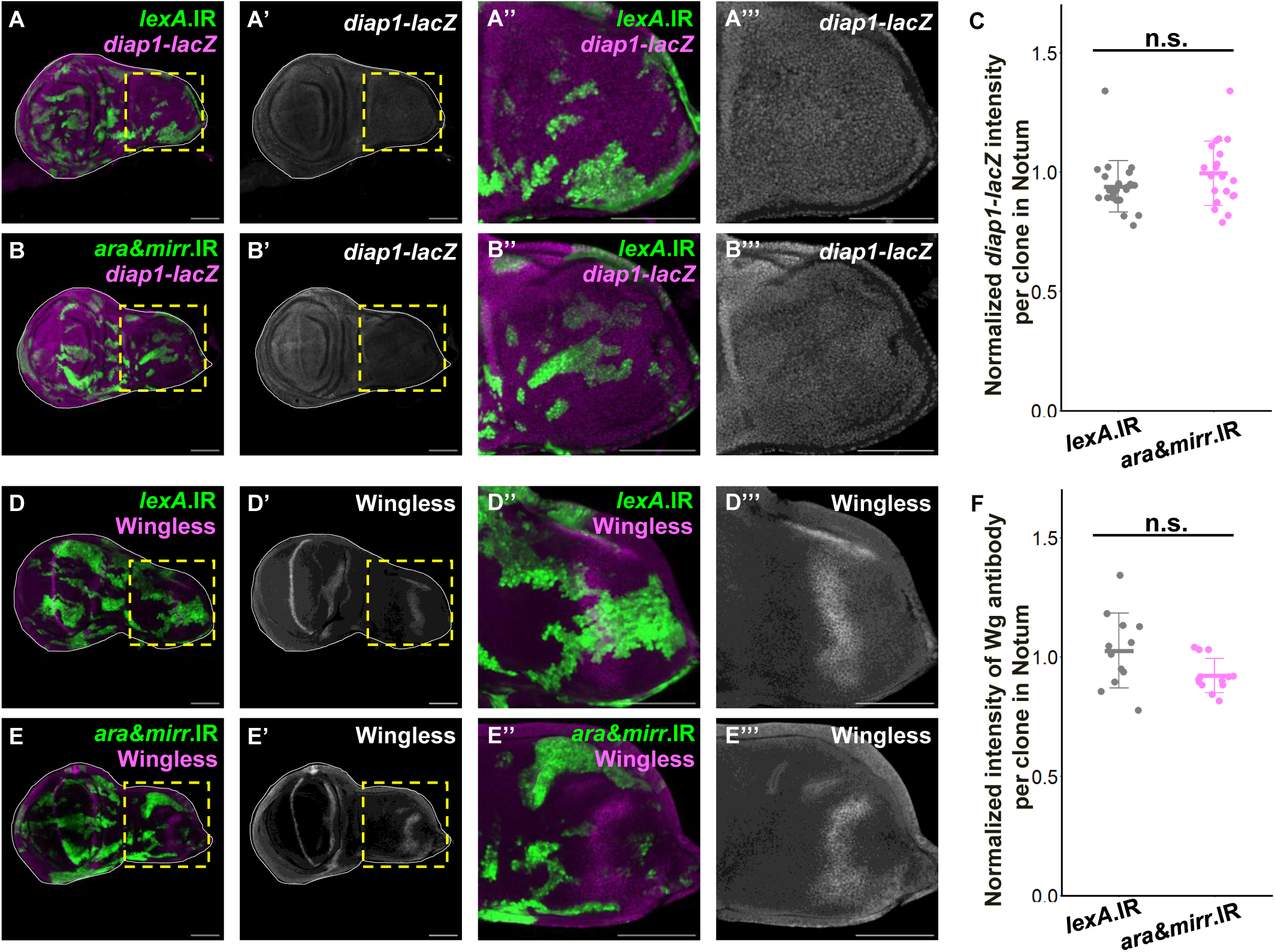
Knockdown of Iro-C does not affect Hippo or Wingless pathway in the notum region. (A and B) Wing disc bearing GFP-labeled *lexA*.IR (A), or *ara&mirr*.IR (B) clones, stained with anti-β-galactosidase antibody for the *diap1-lacZ* reporter, dissected at 96-120 h AEL. (A”, A’’’, B”, and B’’’) Magnified images of the notum region of A, A’, B and B’ respectively. (C) Quantification of the relative intensity of *diap1-lacZ* in *lexA*.IR (n=23) and *ara&mirr.*IR (n=20, p=0.1281) clones. n.s. (not significant); Wilcoxon rank sum test. (D and E) Wing disc bearing GFP-labeled *lexA*.IR (A), or *ara&mirr*.IR (B) clones, stained with Wg antibody, dissected at 96-120 h AEL. (D”, D’’’, E”, and E’’’) Magnified images of the notum region of A, A’, B and B’ respectively. (F) Quantification of the relative intensity of Wg antibody in *lexA*.IR (n=12) and *ara&mirr.*IR (n=13, p=0.0535) clones. n.s. (not significant); Wilcoxon rank sum test. Scale bar, 100 µm.

**Figure S6.**
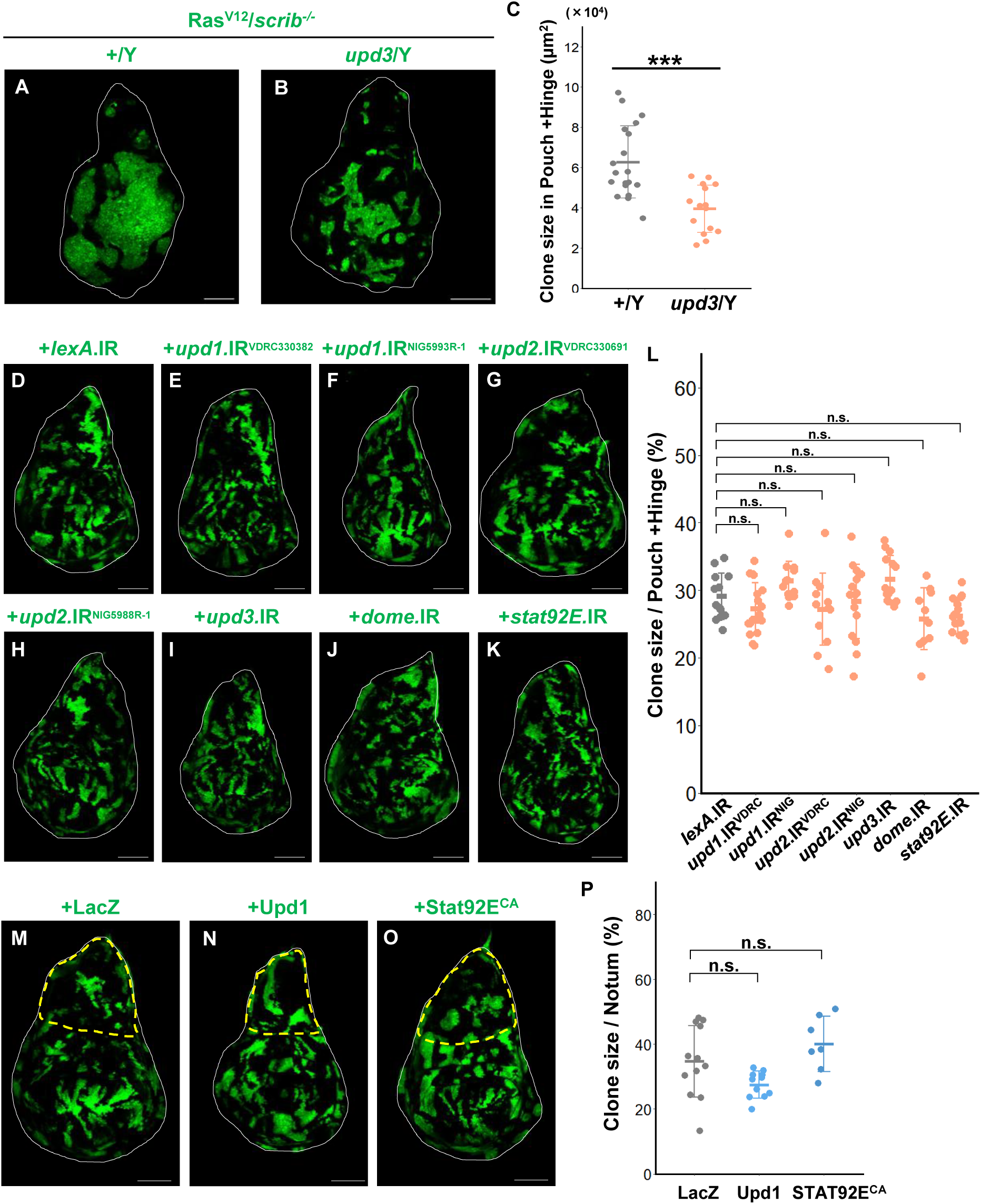
Knockdown or overexpression of Upd-JAK-STAT signaling components in the wild-type background does not affect cell proliferation, Related to Figure 4. (A and B) Wing discs of male larvae bearing GFP-labeled Ras^V12^/*scrib^-/-^*clones in wild-type background (A) and *upd3* mutant background (B), dissected at 120-144 h AEL. (C) Quantification of the clone size in the Pouch+Hinge (P+H) region for (A) (n=19) and (B) (n=15, p=0.0001). ***p < 0.001; Wilcoxon rank sum test. (D-L) Wing discs bearing GFP-labeled *lexA*.IR (D), *upd1.*IR^VDRC330382^ (E), *upd1.*IR^NIG5993R-1^ (F), *upd2.*IR^VDRC330691^ (G), *upd2.*IR^NIG5988R-1^ (H), *upd3.*IR (I), *dome.*IR (J), and *stat92E.*IR (K) clones, dissected at 96-120 h AEL. (L) Quantification of the clone size in the Pouch+Hinge (P+H) region (% of the clone area per region area) for *lexA*.IR (n=13), *upd1.*IR^VDRC330382^ (n=18, p=0.7309), *upd1.*IR^NIG5993R-1^ (n=12, p=0.6557), *upd2.*IR^VDRC330691^ (n=12, p=0.7309), *upd2.*IR^NIG5988R-1^ (n=15, p=0.8222), *upd3.*IR (n=14, p=0.6557), *dome.*IR (n=10, p=0.6557), and *stat92E.*IR (n=15, p=0.3722) clones. n.s. (not significant); Kruskal-Wallis test followed by Dunn’s many-to-one comparison test with Holm correction. (M-O) Wing discs bearing GFP-labeled LacZ (M), Upd1 (N), and Stat92E^CA^ (O) expressing clones, dissected at 96-120 h AEL. (P) Quantification of the clone size in the notum region (% of the clone area per region area) for LacZ (n=12), Upd1 (n=11, p=0.1068), and Stat92E^CA^ (n=7, p=0.2247) expressing clones. n.s. (not significant); Kruskal-Wallis test followed by Dunn’s many-to-one comparison test with Holm correction. Scale bar, 100 µm.

**Figure S7.**
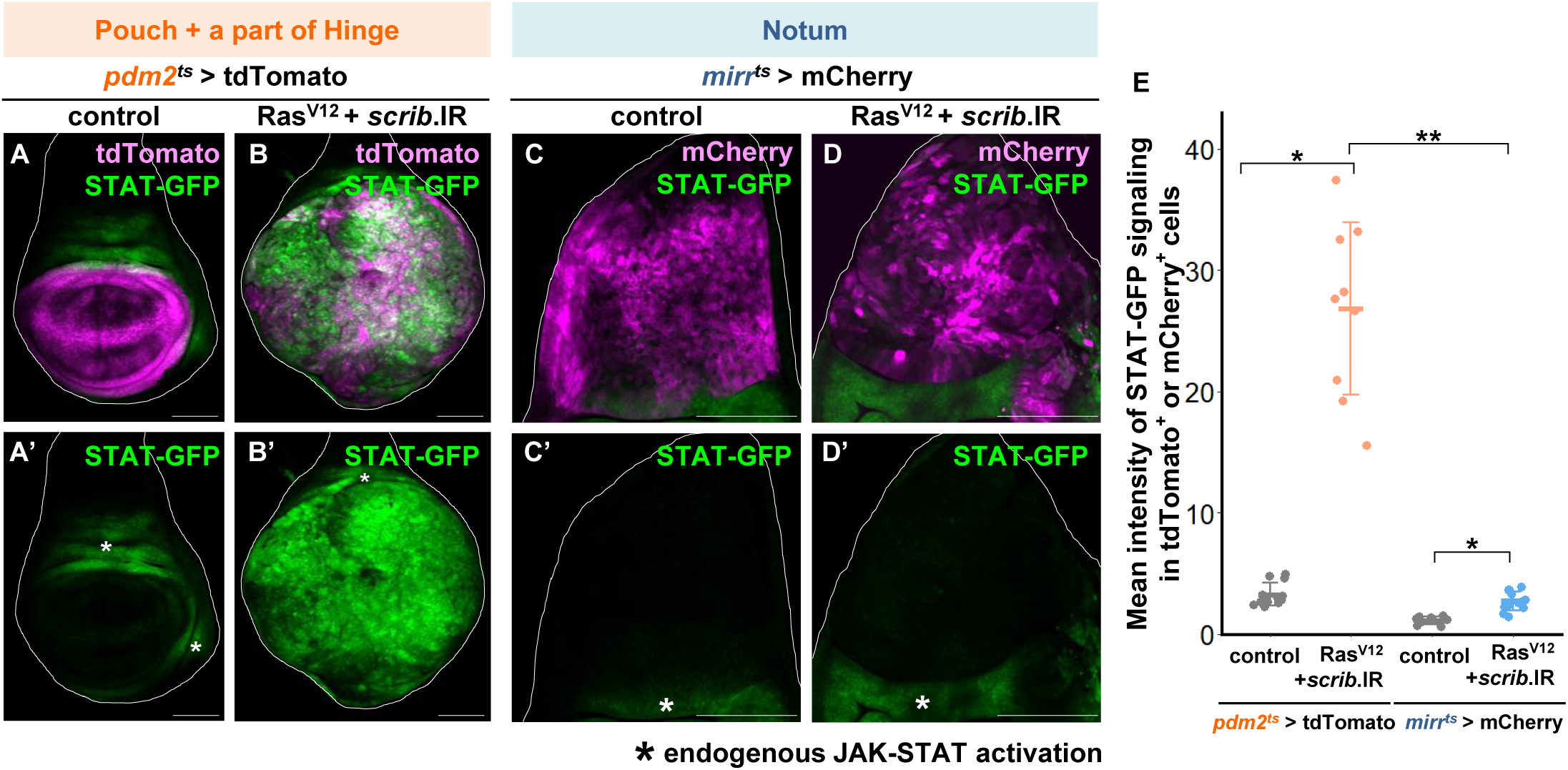
JAK-STAT signaling is not activated in Ras^V12^*/scrib*.IR cells induced in the notum region, Related to Figure 4. (A, B) STAT-GFP expression in a wing disc bearing tdTomato-labeled control and Ras^V12^/*scrib*.IR cells induced by *pdm2*-Gal4. (C, D) STAT-GFP expression in a wing disc bearing mCherry-labeled control and Ras^V12^/*scrib*.IR cells induced by *mirr*-Gal4. (E) Quantification of the mean pixel intensity of STAT-GFP in the tdTomato- or mCherry-labeled area for *pdm2*^ts^>tdTomato (n=12), *pdm2*^ts^>tdTomato, Ras^V12^+*scrib*.IR (n=9,), *mirr*^ts^>mCherry (n=9), and *mirr*^ts^>mCherry, Ras^V12^+*scrib*.IR (n=12). *pdm2*^ts^>tdTomato vs. *pdm2*^ts^>tdTomato, Ras^V12^+*scrib*.IR, p=0.0201; *mirr*^ts^>mCherry vs. *mirr*^ts^>mCherry, Ras^V12^+*scrib*.IR, p=0.0201; *pdm2*^ts^>tdTomato, Ras^V12^+*scrib*.IR vs. *mirr*^ts^>mCherry, Ras^V12^+*scrib*.IR, p=0.0031.**p < 0.01; *p < 0.05; Kruskal-Wallis test followed by Dunn’s all-pairs comparison test with Holm correction.

**Figure S8.**
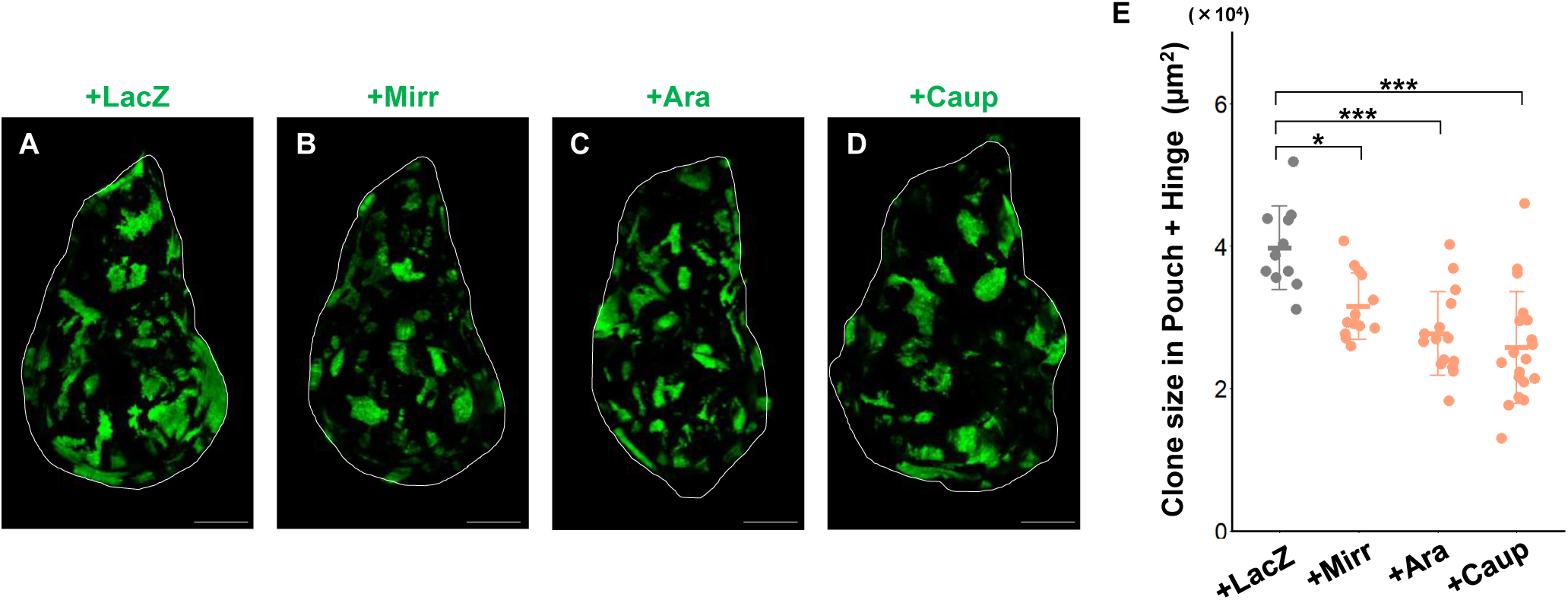
Forced expression of Iro-C only mildly affects wild-type cell proliferation, Related to Figure 5. (A-D) Wing discs bearing GFP-labeled LacZ (A), Mirr (B), Ara (C), and Caup (D) expressing clones, dissected at 96-120 h AEL. (E) Quantification of the clone size in the Pouch+Hinge (P+H) region for LacZ (n=11), Mirr (n=13, p=0.0398), Ara (n=15, p=0.0005), and Caup (n=19, p<0.0001) expressing clones. ***p < 0.001; n.s. (not significant); Kruskal-Wallis test followed by Dunn’s many-to-one comparison test with Holm correction.

**Figure S9.**
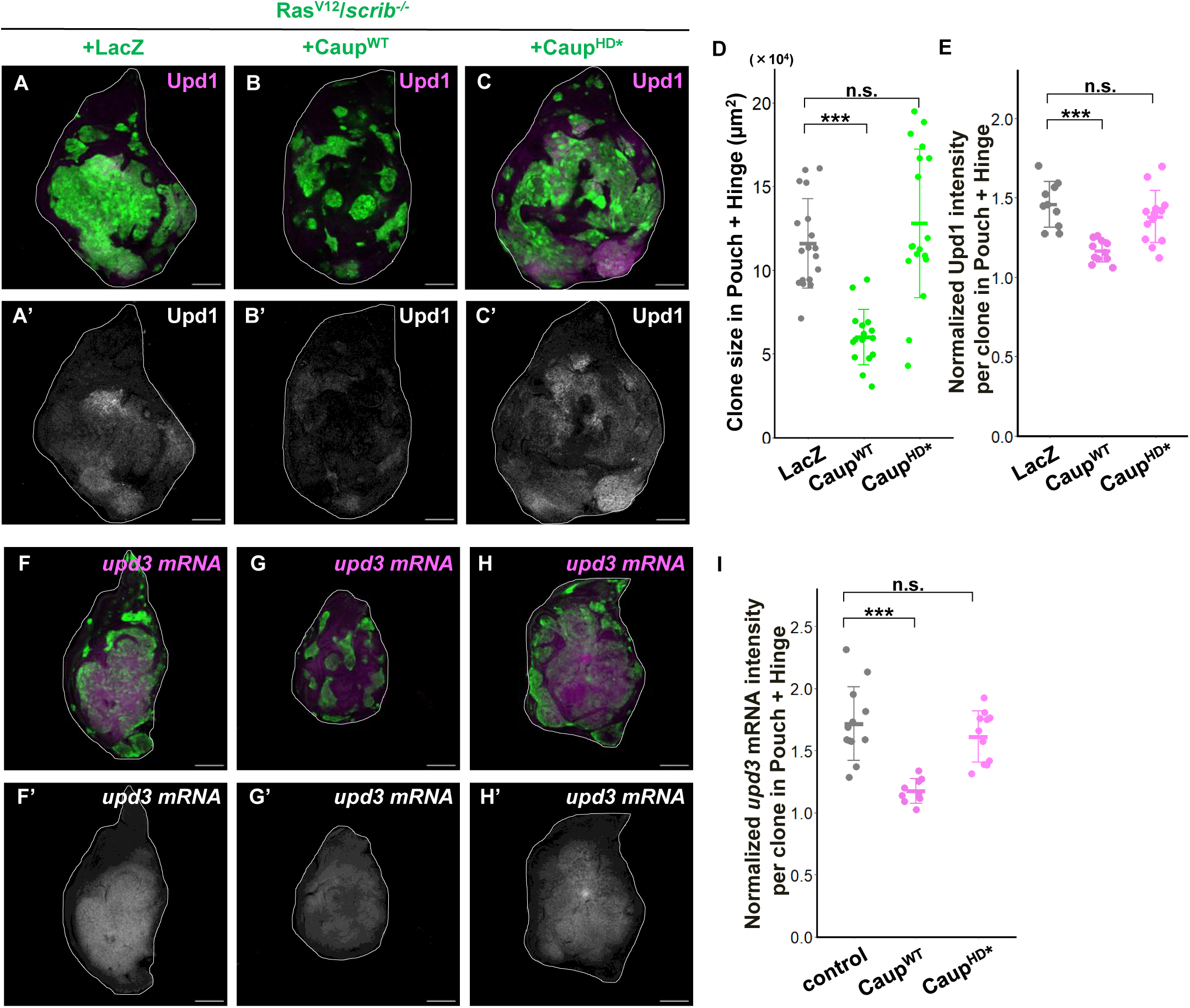
Forced expression of Iro-C suppresses tumorigenesis in the tumor-prone regions, Related to Figure 5. (A-C) Wing discs bearing GFP-labeled Ras^V12^/*scrib^-/-^*+LacZ (A), Ras^V12^/*scrib^-/-^*+Caup^WT^ (B), and Ras^V12^/*scrib^-/-^*+Caup^HD*^ (C) expressing clones stained with anti-Upd1 antibody, dissected at dissected at 144-168 h AEL. (D) Quantification of the clone size in the Pouch+Hinge (P+H) region for Ras^V12^/*scrib^-/-^*+LacZ (n=18), Ras^V12^/*scrib^-/-^*+Caup^WT^ (n=16, p<0.0001), and Ras^V12^/*scrib^-/-^*+Caup^HD*^ (n=19, p=0.4567) expressing clones. ***p < 0.001; n.s. (not significant); Kruskal-Wallis test followed by Dunn’s many-to-one comparison test with Holm correction. (E) Quantification of the relative intensity of Upd1 antibody for Ras^V12^/*scrib^-/-^*+LacZ (n=10), Ras^V12^/*scrib^-/-^*+Caup^WT^ (n=11, p=0.0001), and Ras^V12^/*scrib^-/-^*+Caup^HD*^ (n=13, p=0.2377) expressing clones. ***p < 0.001; n.s. (not significant); Kruskal-Wallis test followed by Dunn’s many-to-one comparison test with Holm correction. (F-H) Wing discs bearing GFP-labeled Ras^V12^/*scrib^-/-^*+LacZ (F), Ras^V12^/*scrib^-/-^*+Caup^WT^ (G), and Ras^V12^/*scrib^-/-^*+Caup^HD*^ (H) expressing clones hybridized with *upd3* RNA probe to visualize *upd3 mRNA* expression, dissected at 144-168 h AEL. (I) Quantification of the relative intensity of *upd3* FISH signal in Ras^V12^/*scrib^-/-^*+LacZ (n=12), Ras^V12^/*scrib^-/-^*+Caup^WT^ (n=9, p=0.0001), and Ras^V12^/*scrib^-/-^*+Caup^HD*^ (n=11, p=0.6549) expressing clones. ***p < 0.001; n.s. (not significant); Kruskal-Wallis test followed by Dunn’s many-to-one comparison test with Holm correction.

**Figure S10.**
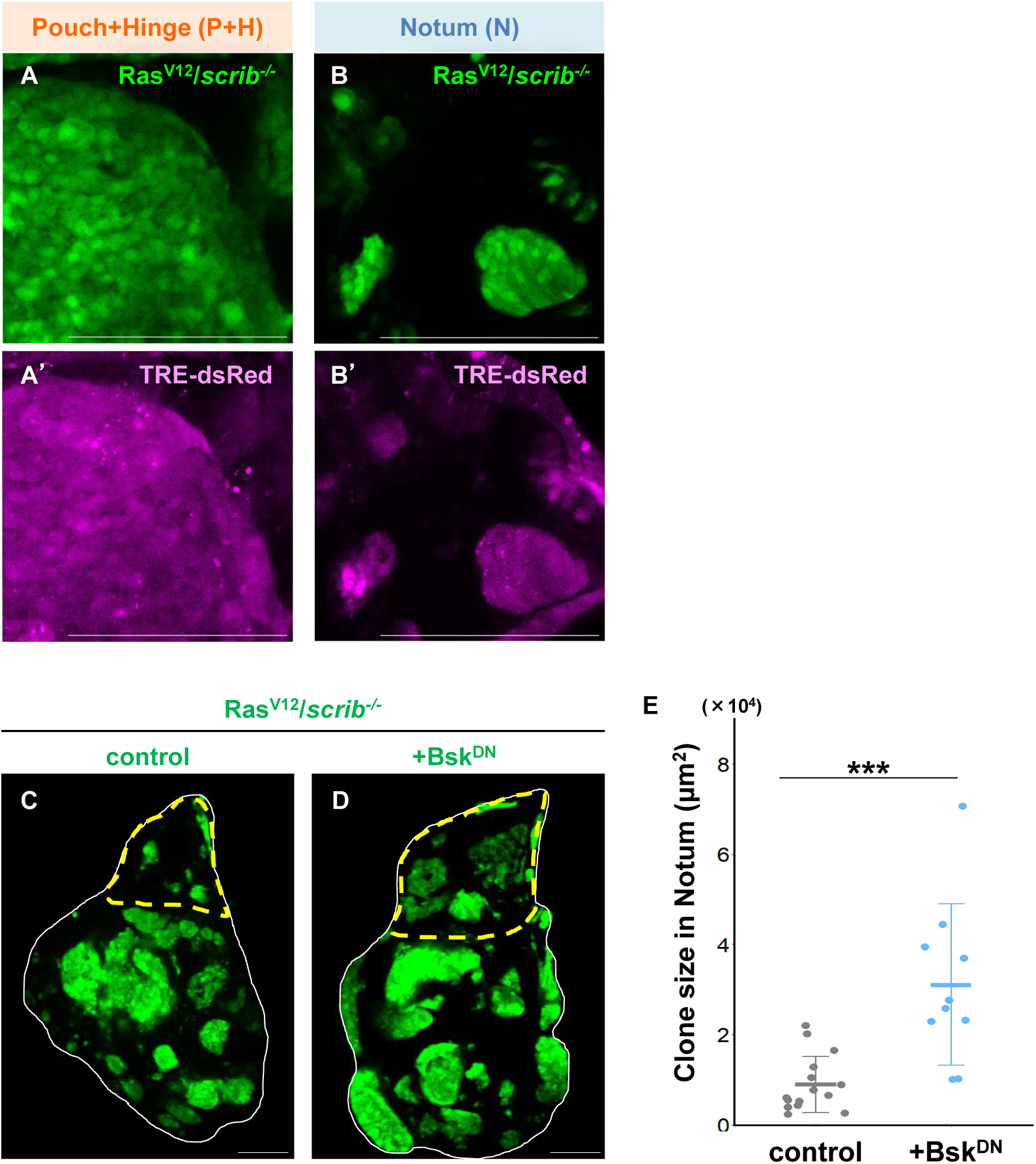
Elevated JNK signaling exerts an anti-growth effect on RasV12/scrib-/- tumorsin the notum region. (A, B) TRE-dsRed expression in a wing disc bearing GFP-labeled Ras^V12^/*scrib^-/-^* cells within the Pouch+Hinge (P+H) region (A) and Notum (N) region (B). (C, D) Wing discs bearing GFP-labeled Ras^V12^/*scrib^-/-^* cells (C), and Ras^V12^/*scrib^-/-^*+Bsk^DN^-expressing cells (D). (E) Quantification of the clone size in the notum region for Ras^V12^/*scrib^-/-^* cells (n=15), and Ras^V12^/*scrib^-/-^*+Bsk^DN^-expressing cells (n=10, p<0.0001). ***p < 0.001; Wilcoxon rank sum test. Scale bar, 100 µm.

**Table 1.**
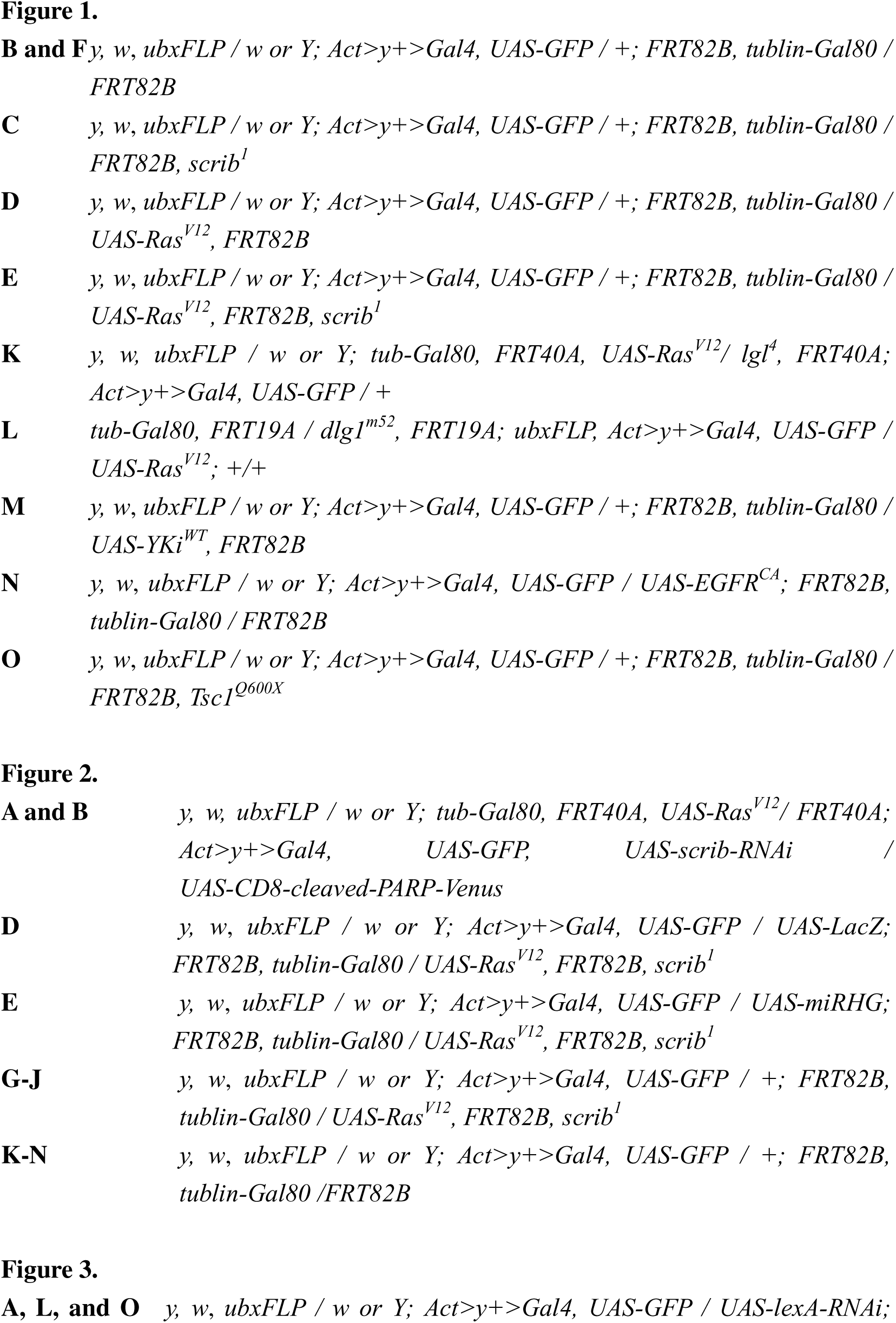
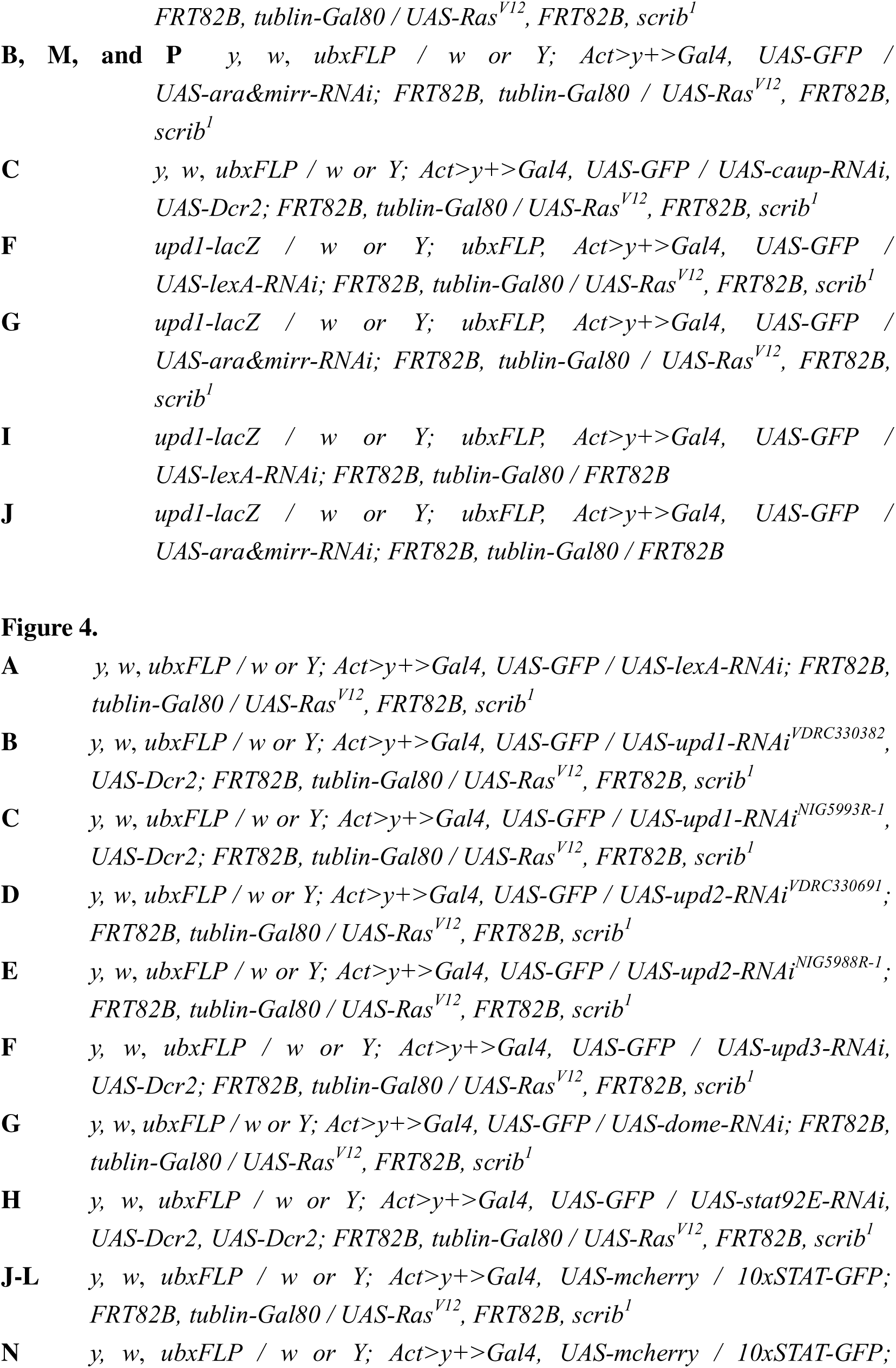
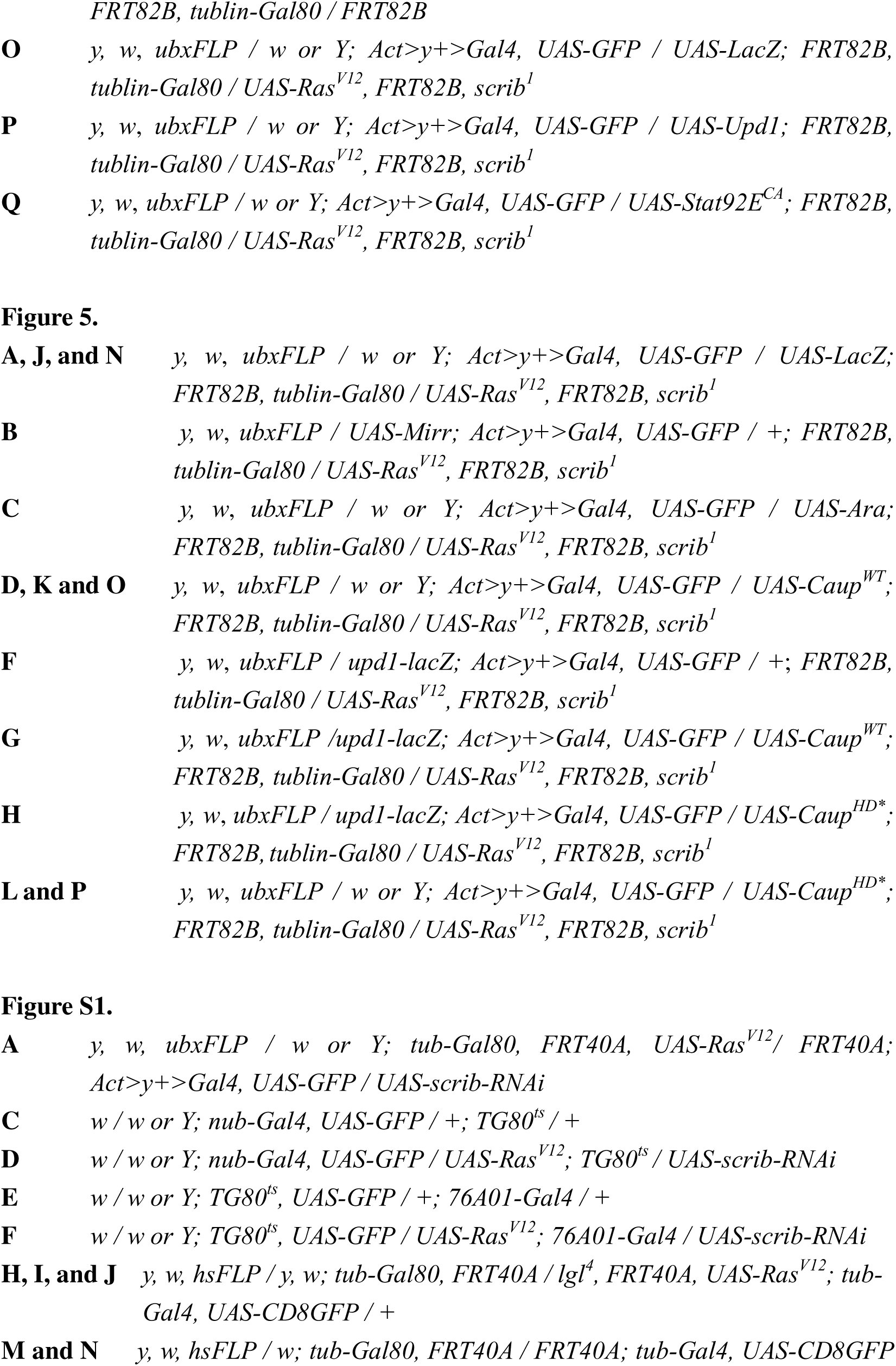
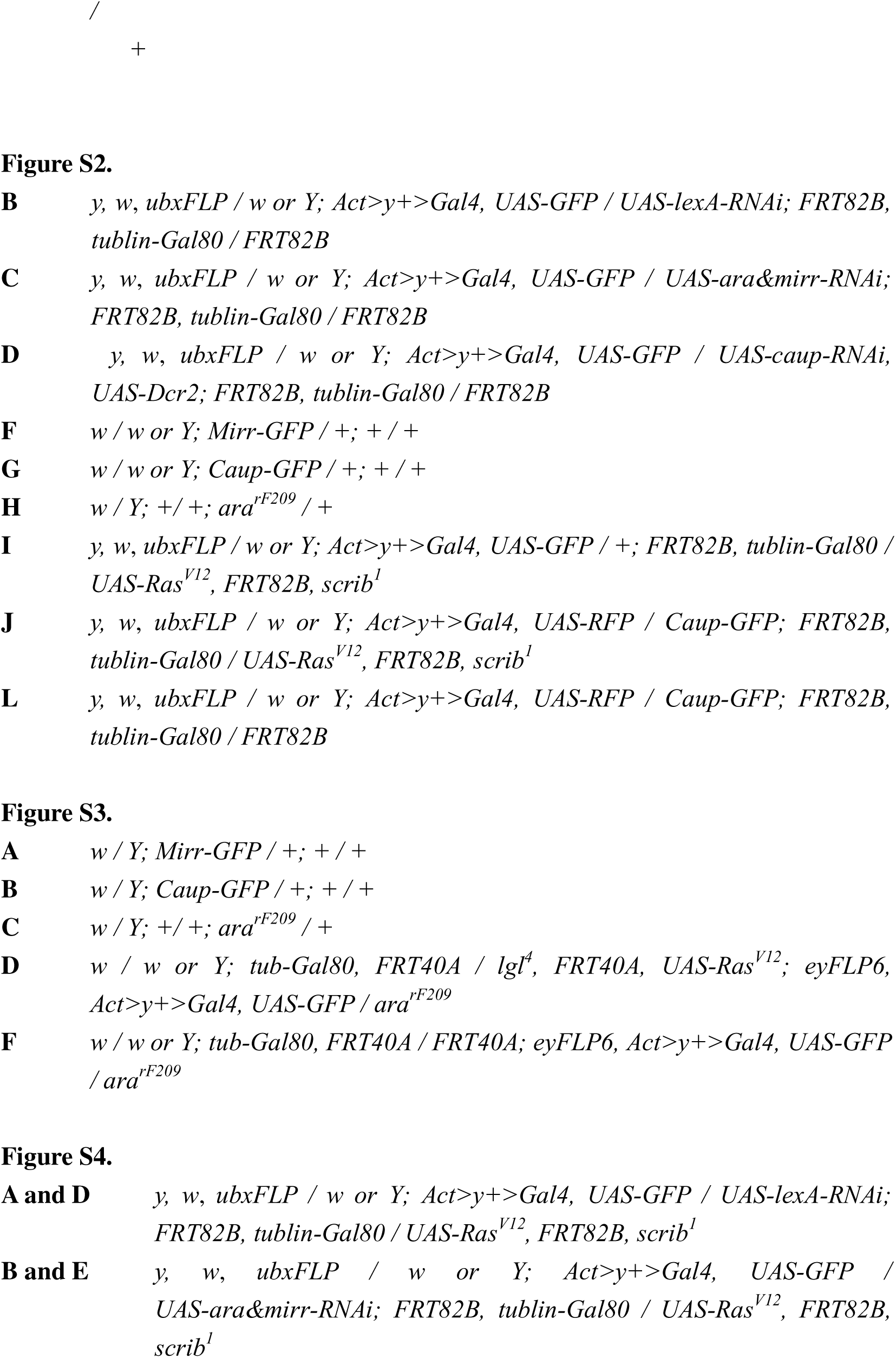
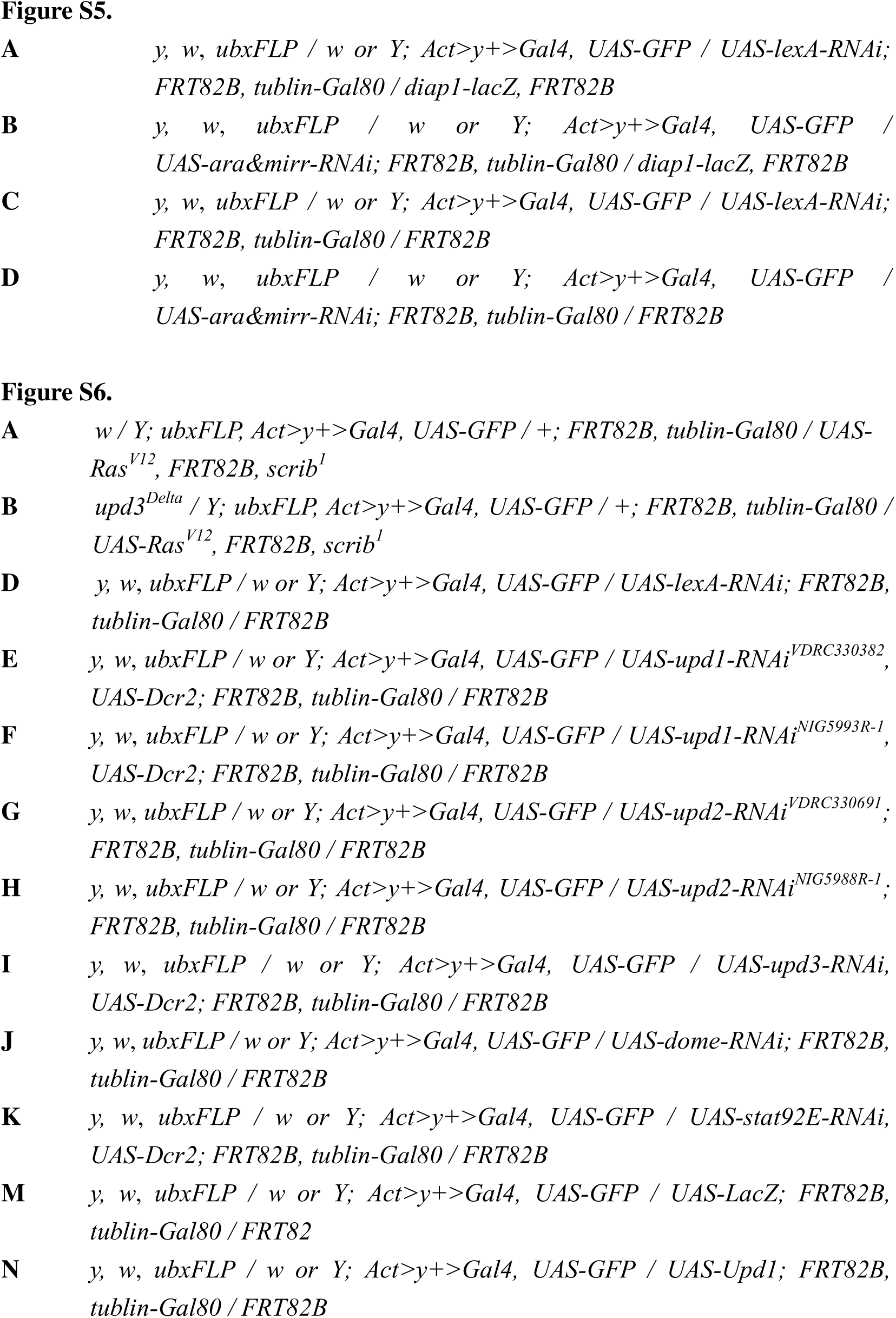
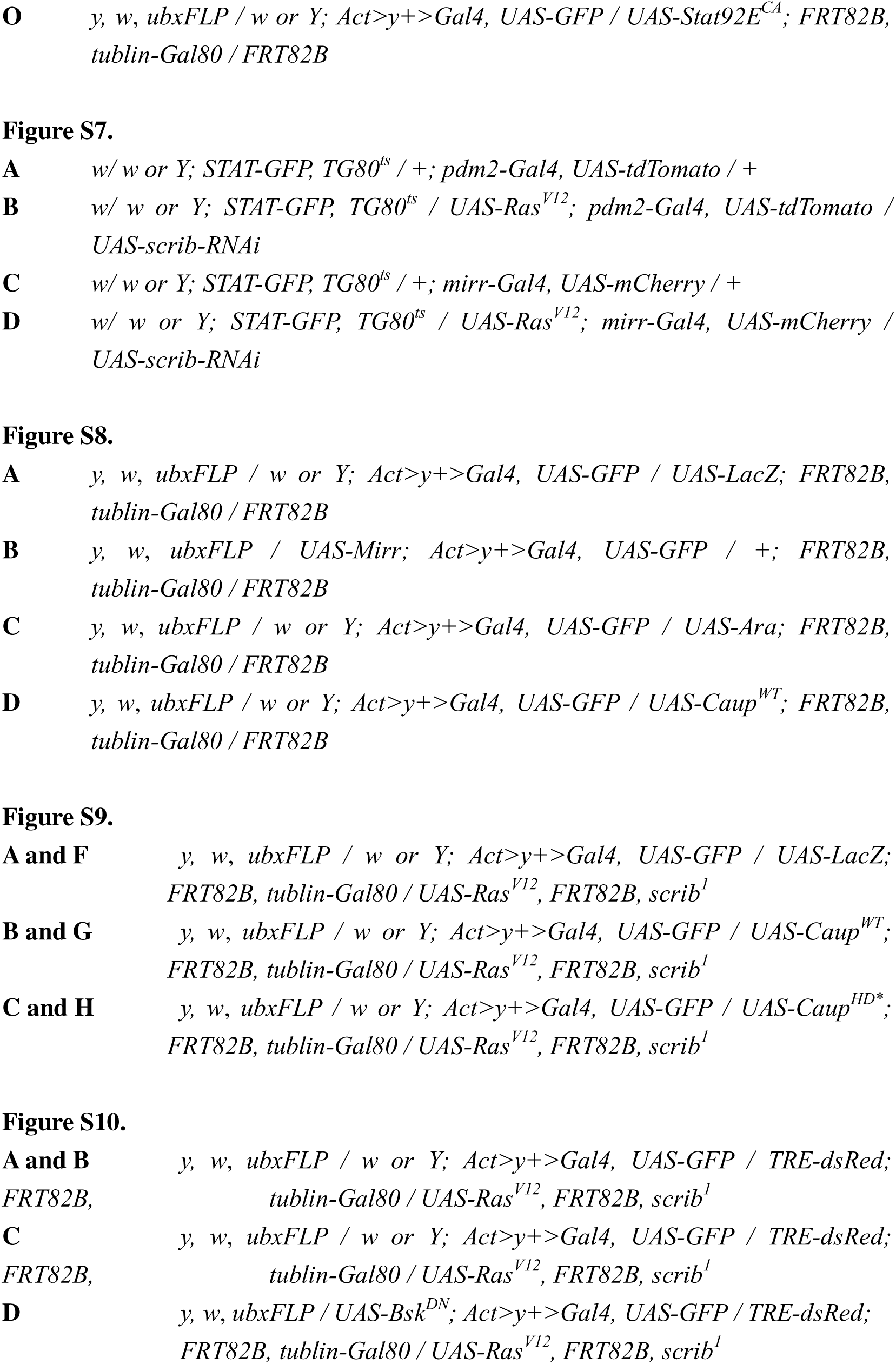
Detailed Genotypes used in each figure, Related to Figure 1-5 and S1- S7.

## Reagents and tools table

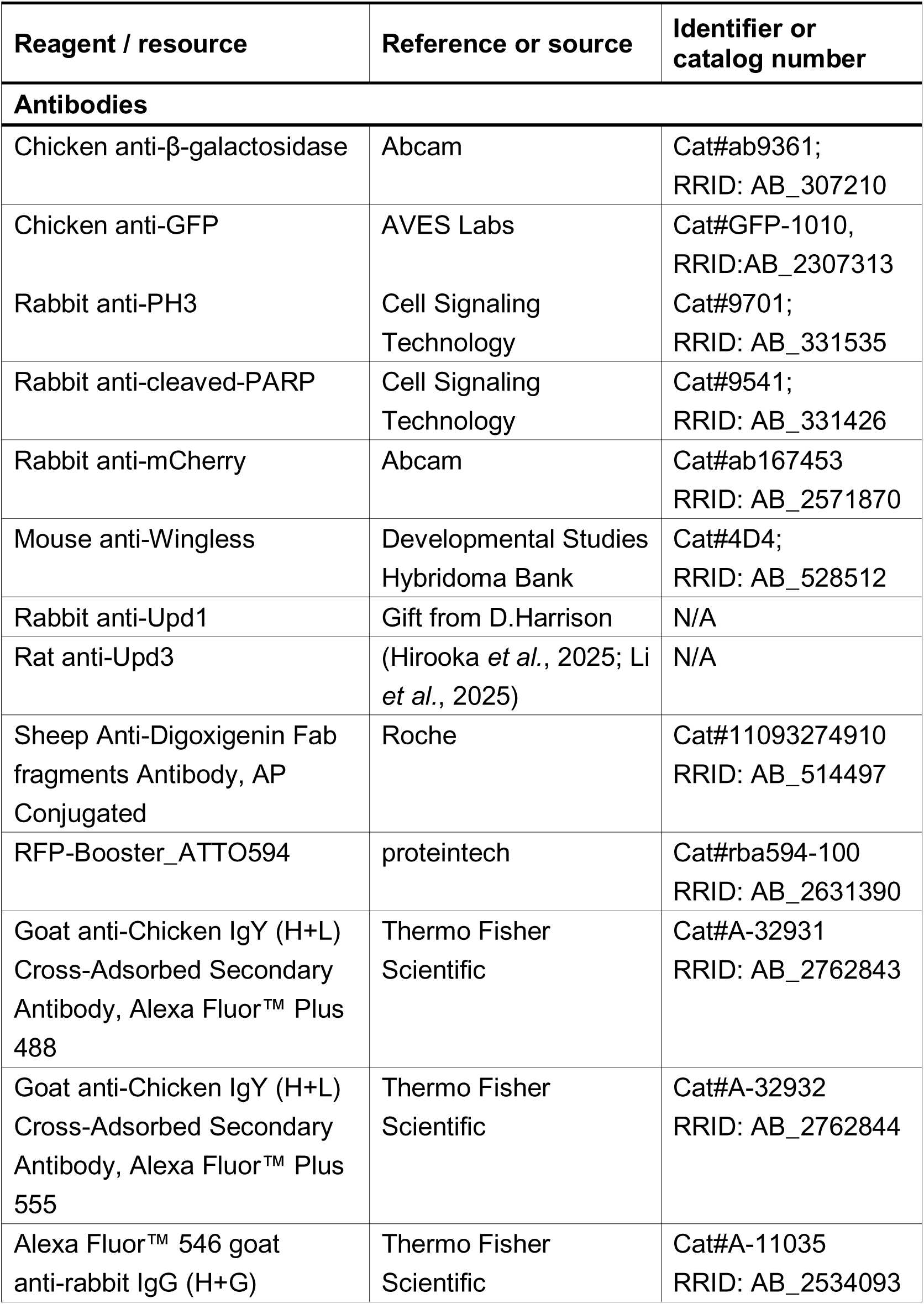
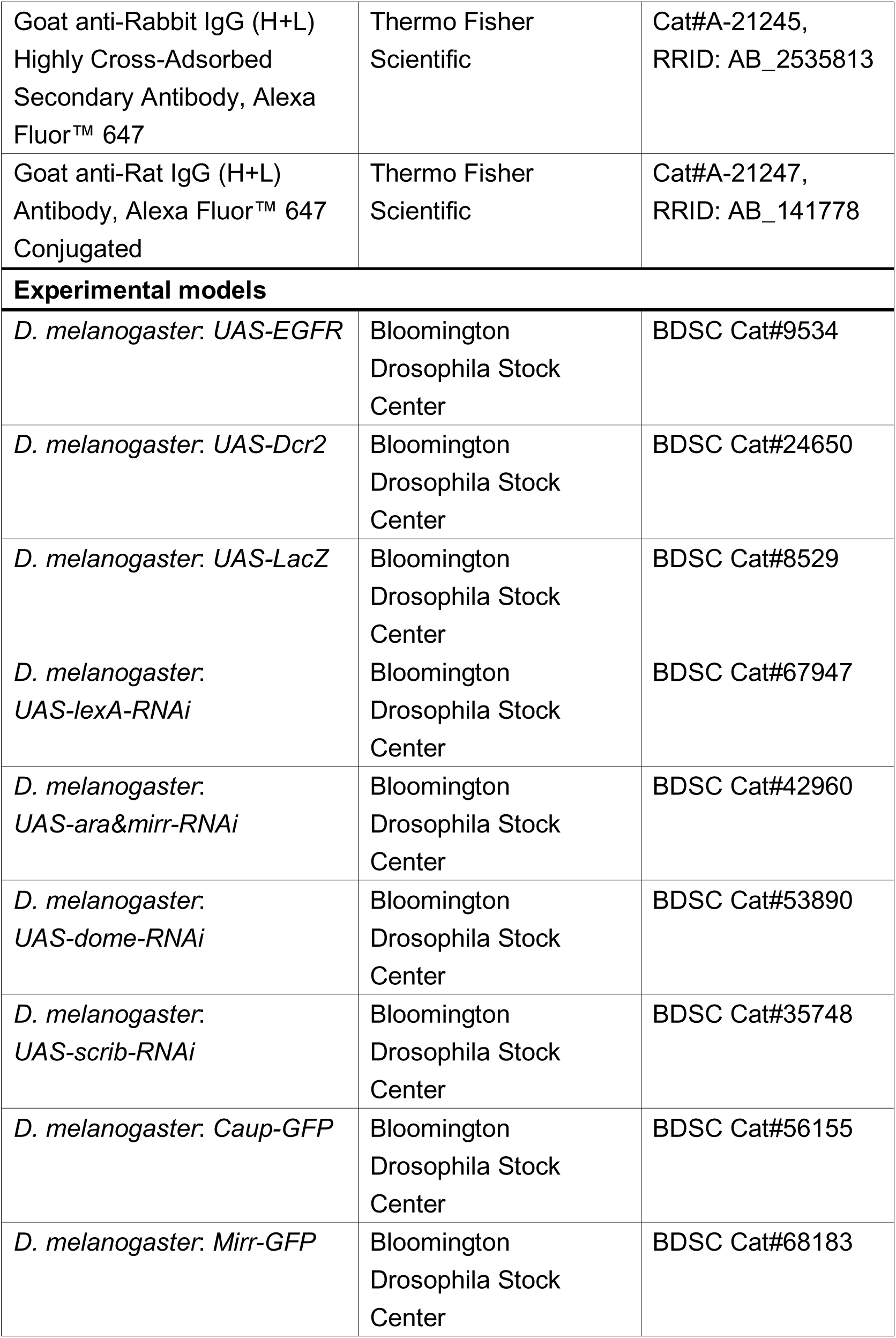
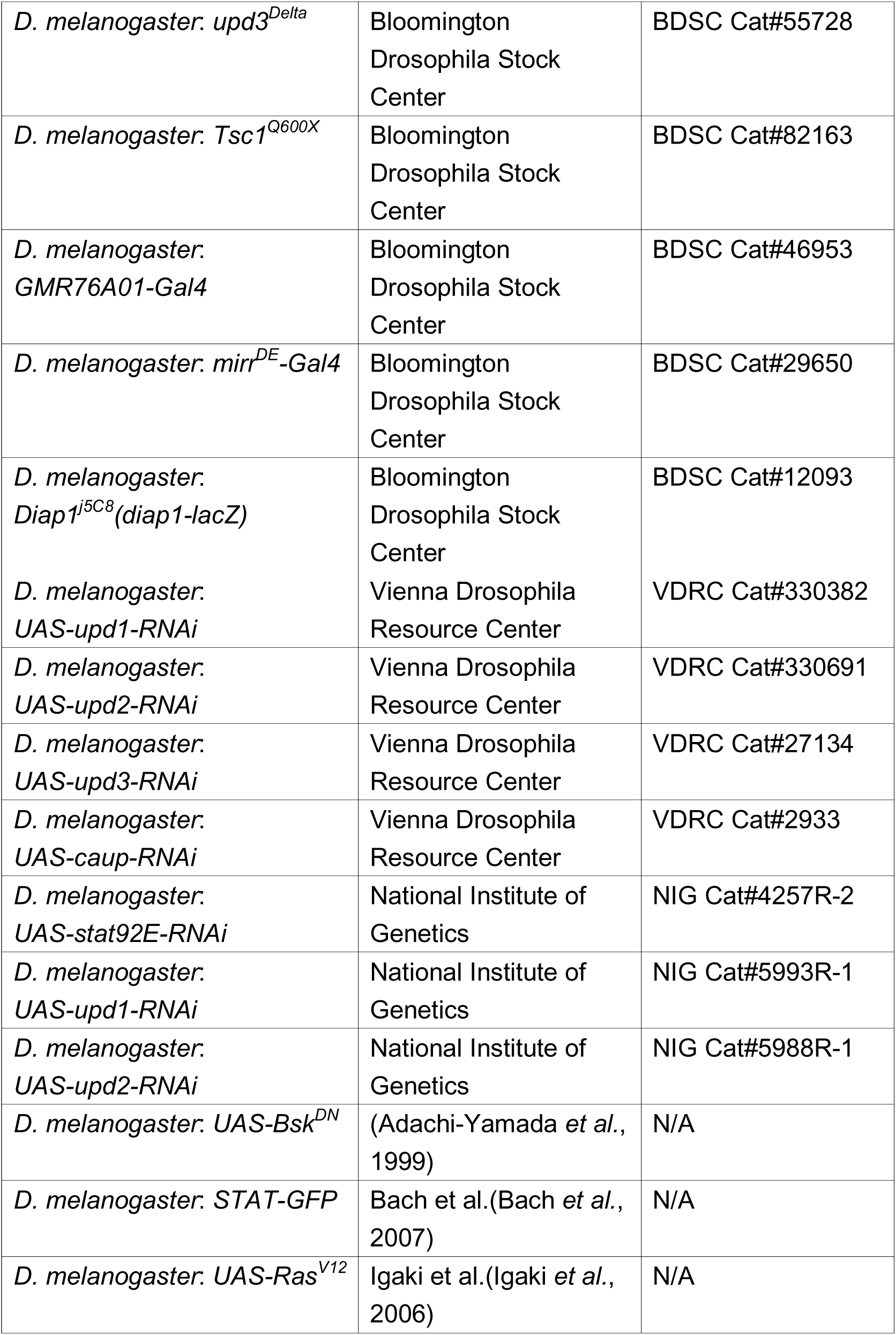
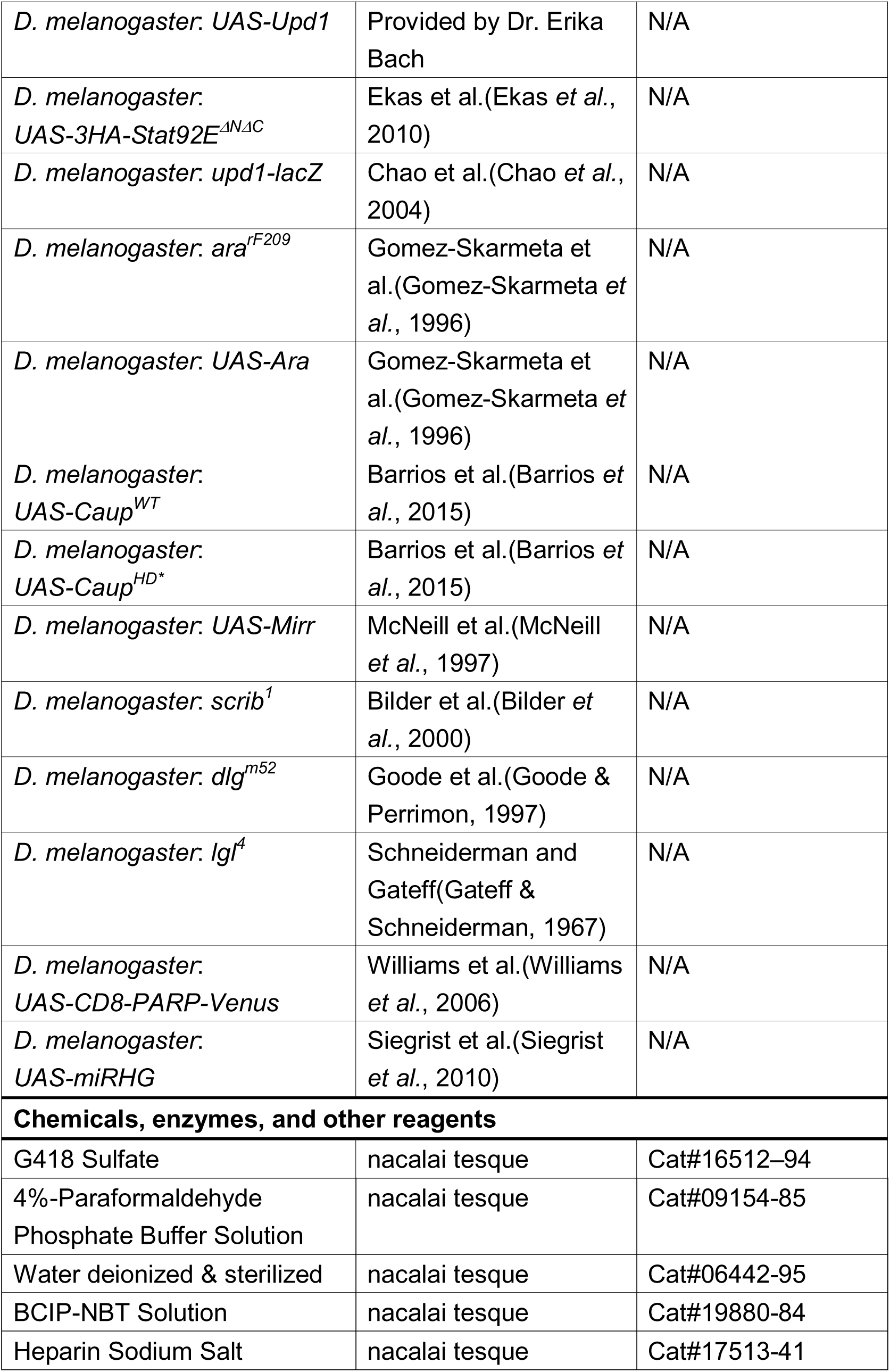
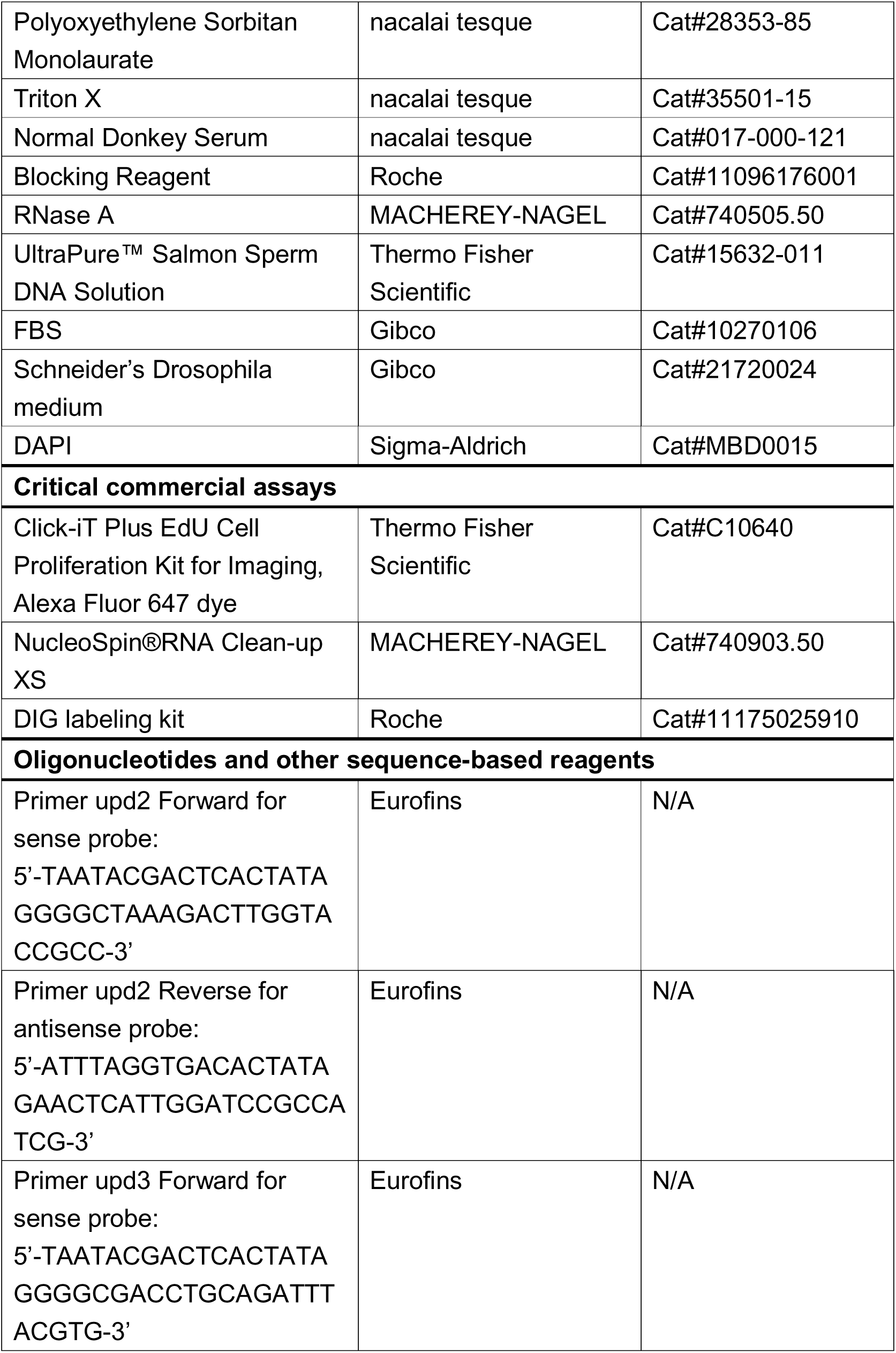
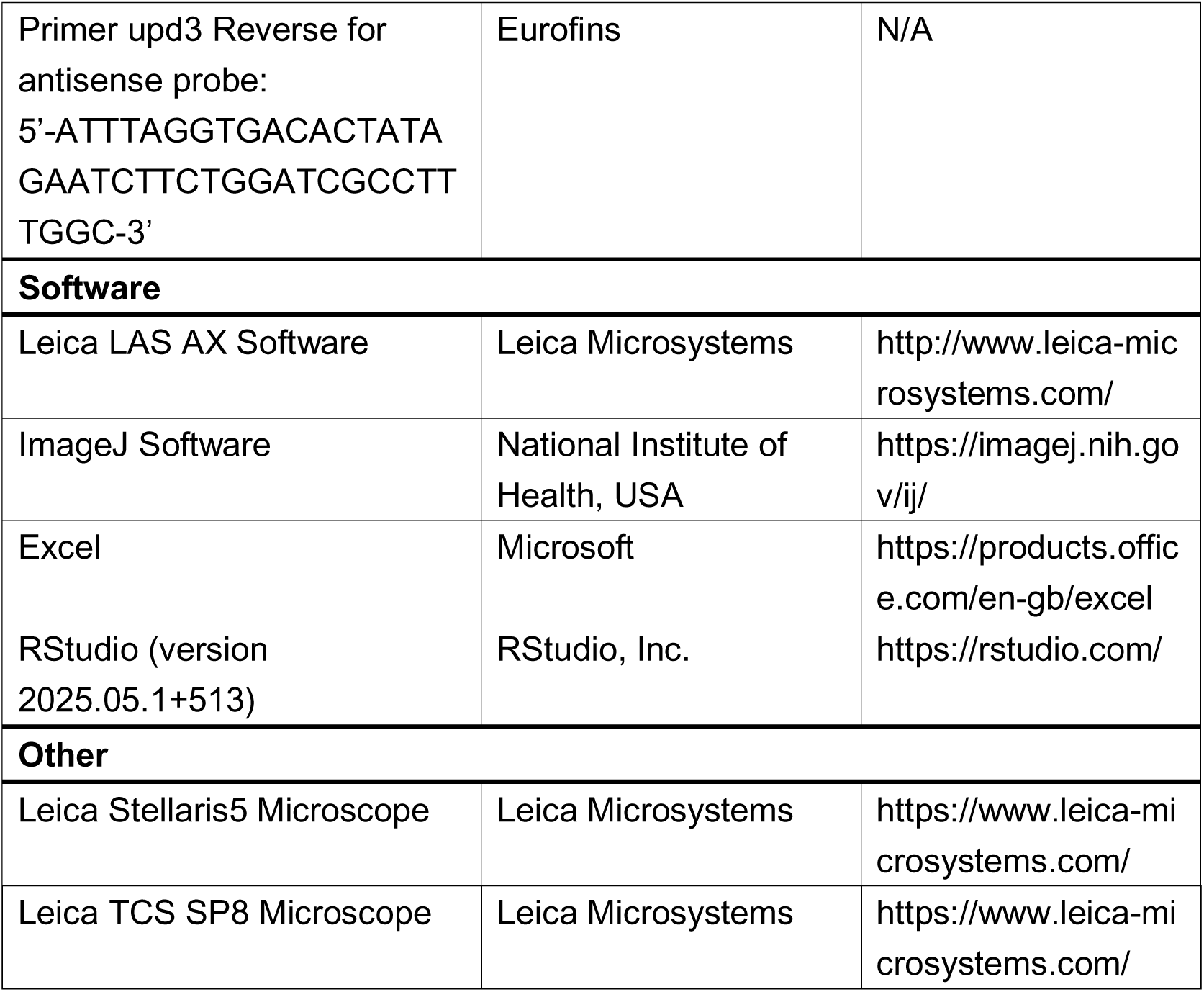

